# Asymmetric learning facilitates human inference of transitive relations

**DOI:** 10.1101/2021.04.03.437766

**Authors:** Simon Ciranka, Juan Linde-Domingo, Ivan Padezhki, Clara Wicharz, Charley M. Wu, Bernhard Spitzer

## Abstract

Humans and other animals are capable of inferring never-experienced relations (e.g., A>C) from other relational observations (e.g., A>B and B>C). The processes behind such transitive inference are subject to intense research. Here, we demonstrate a new aspect of relational learning, building on previous evidence that transitive inference can be accomplished through simple reinforcement learning mechanisms. We show in simulations that inference of novel relations benefits from an asymmetric learning policy, where observers update only their belief about the winner (or loser) in a pair. Across 4 experiments (n=145), we find substantial empirical support for such asymmetries in inferential learning. The learning policy favoured by our simulations and experiments gives rise to a compression of values which is routinely observed in psychophysics and behavioural economics. In other words, a seemingly biased learning strategy that yields well-known cognitive distortions can be beneficial for transitive inferential judgments.

## Main

Humans routinely infer relational structure from local comparisons. For instance, learning that boxer Muhammad Ali defeated George Foreman can let us infer that Ali would likely win against other boxers that Foreman had defeated. More formally, generalizing from relational observations to new, unobserved relations (e.g., knowing A>B and B>C leads to A>C) is commonly referred to as transitive inference^1–4^. Transitive inference is not a uniquely human capacity ^5^ but can also be observed in non-human primates ^6–8^, rats ^9^, and birds ^10–12^.

In the laboratory, transitive inference can be observed after teaching subjects the relations between neighbouring elements from an ordered set of arbitrary stimuli (**Fig. 1a**). The neighbour relations are typically taught through pairwise choice feedback (**Fig. 1b**) where the relational information is deterministic (i.e., if A>B, in our sporting analogy, A would never lose a match against B). Various theories have been proposed to describe how observers accomplish transitive inferences of non-neighbour relations (e.g., A>D) in such settings. One class of models posits that subjects learn implicit value representations for each individual element (A, B, C, etc.), which then enables judgments of arbitrary pairings ^3,13,14^. Alternatively, transitive inference could be accomplished through more explicit, hippocampus-based memory processes ^15–18^, which we will return to further below.

**Fig. 1.**
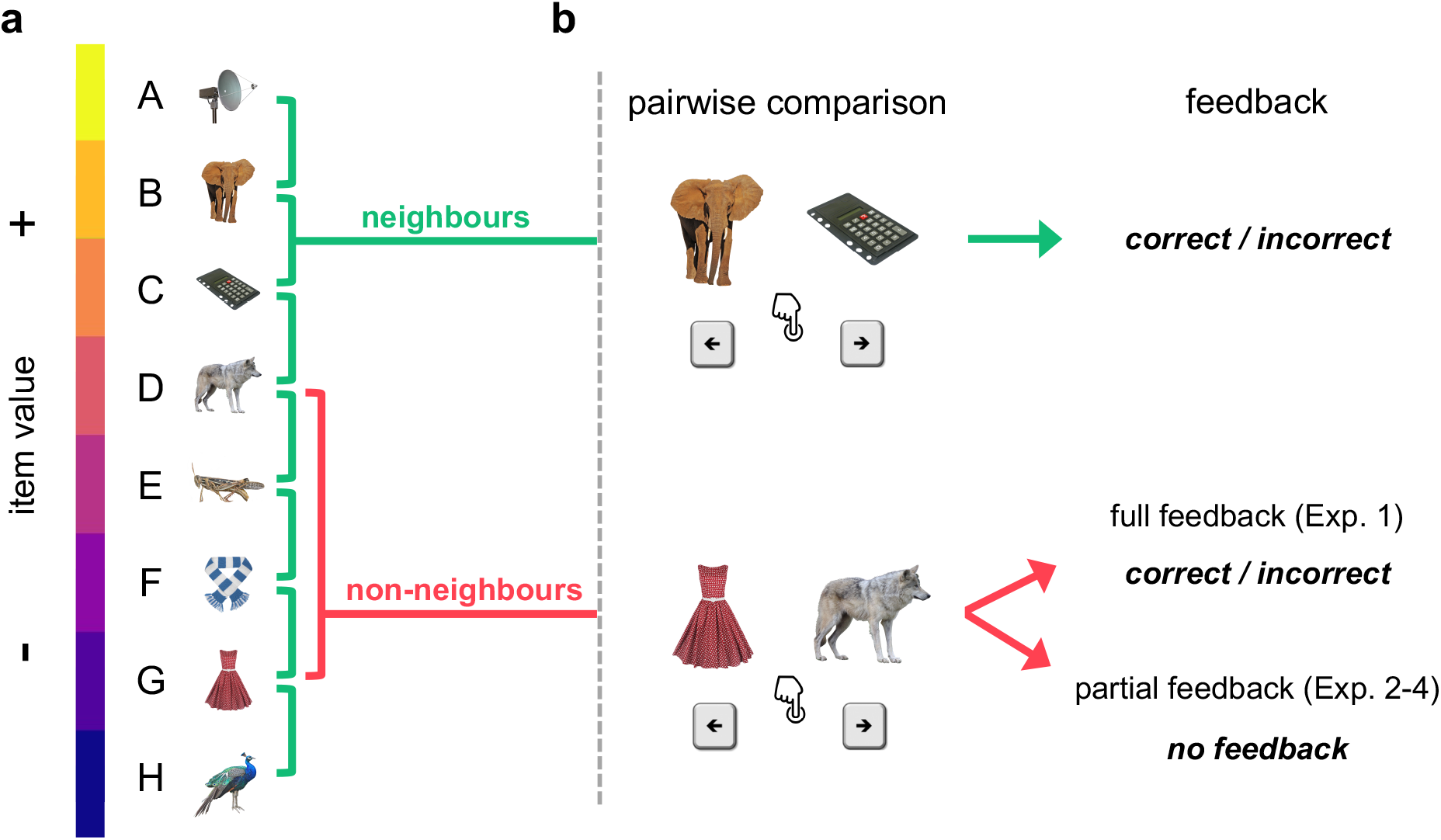
Relational learning paradigm. **a**, Exemplary stimulus set and hidden relational value structure. **b**, Example trials for pairwise comparisons of neighbouring (*top*) and non-neighbouring items (*bottom*). Participants are asked on each trial to select the higher-valued item. Choices on neighbour trials are always given feedback. Choices on non-neighbour trials are given feedback in the full-feedback condition, but not in the partial feedback condition (see text for details).

Before turning to transitive inference, we consider relational learning in a “full feedback” scenario (cf. Fig. 1b) where choice feedback is provided for every possible pairing of items, such that no transitive inference is required. We model implicit value learning in this setting through a simple reinforcement learning (RL) mechanism (*Q*-learning, see Methods) by which relational feedback (e.g., “correct” when selecting A over B) may increase the perceived value (*Q*) of item A and decrease that of item B (Model **Q1, Fig. 2a**). In this simple RL model, relational feedback *symmetrically* updates (with opposite signs) the value estimates for both items in a pair. For instance, if Muhammad Ali beat George Foreman, it seems rational to attribute this outcome to Ali’s greater skill as much as to Foreman’s deficit. We show in simulations that symmetric value updating is in fact optimal in the full feedback setting. An alternative model with *asymmetric* learning rates (α^+^ ≠ α^-^) applied to the winner and loser in a pair, respectively (Model **Q2**; “2” denotes dual learning rates), learns worse than the symmetric model (Q1) where *α*^+^ = *α*^−^ (**Fig. 2b-c**). Implicit value learning generally gives rise to a “symbolic distance effect” ^1,19,20^, where nearby elements are less discriminable (due to more similar value estimates) than elements with greater ordinal distance ^14,21^.

**Fig. 2.**
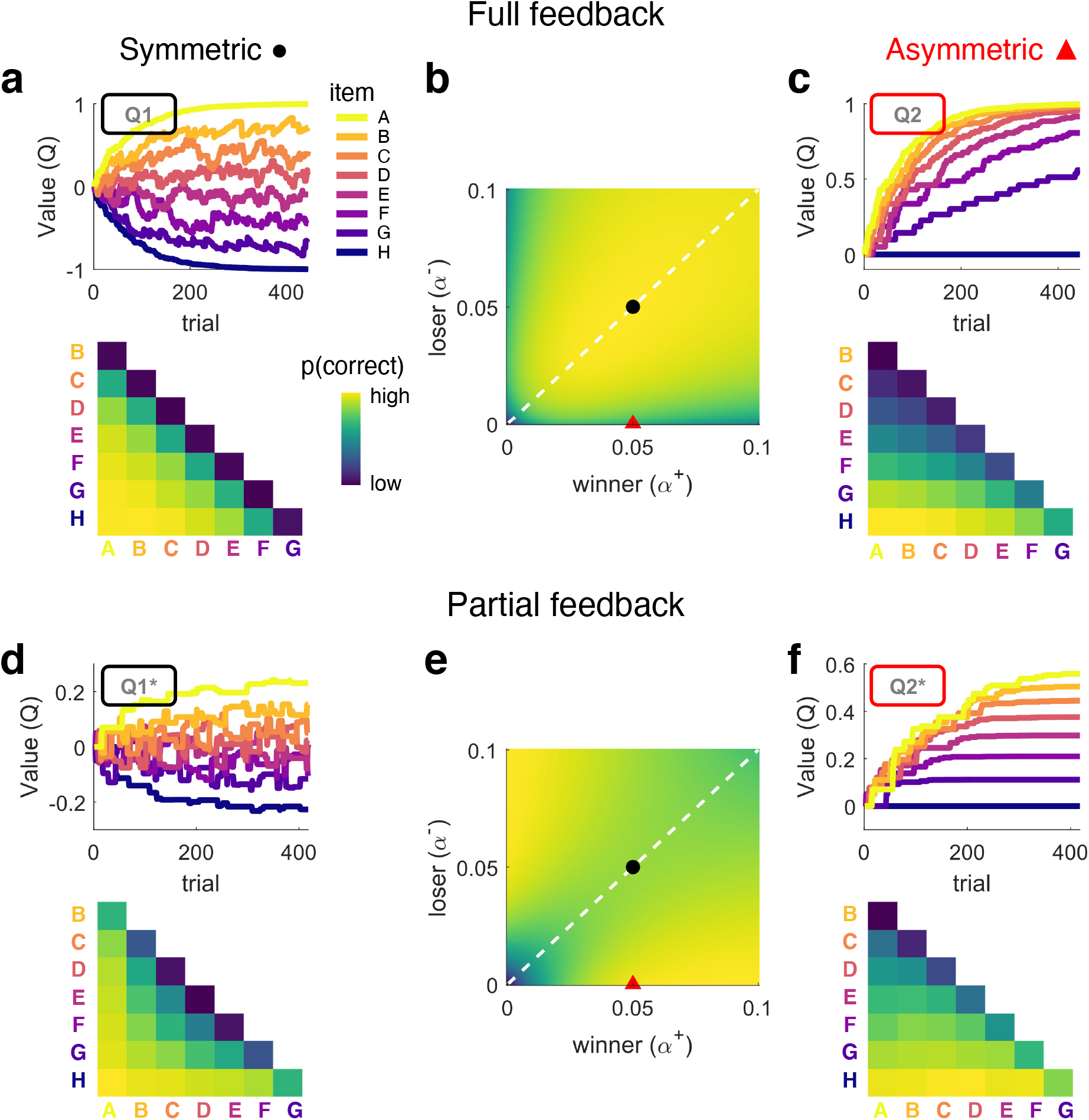
Model simulations under full (a-c) and partial feedback (d-f). **a**, Item-level learning under full feedback (see Exp. 1) simulated with symmetric model Q1. *Top*, exemplary evolution of item values *Q* (a.u.) over trials. *Bottom*, simulated probability of making a correct choice for each item pairing (aggregated across all trials in the top panel). **b**, Simulated task performance (mean proportion correct choices on the second half of trials) of asymmetric model Q2 across different learning rates *α*^+^(winning items) and *α*^−^(losing items). For values on the diagonal (dashed white line), Q2 is equivalent to Q1. Black dot indicates parameters used for simulation of symmetric learning in panel *a*. Red triangle indicates parameters used for simulation of asymmetric learning in panel *c*. **c**, Same as *a*, but using model Q2 with asymmetric learning rates. **d**, Same as *a*, but for model Q1* in a partial feedback scenario (see Exps. 2-4). **e and f**, Same as *b* and *c*, but using model Q2* under partial feedback. Note that asymmetric learning leads to lower performance under full feedback (*b*) but improves performance under partial feedback (*e*). Asymmetric learning results in a compressed value structure that is asymptotically stable under partial feedback (*f*) but not under full feedback (*c*).

Next, we turn to a “partial feedback” setting, which is the typical transitive inference scenario, with feedback only being provided for pairs of items with neighbouring values (Fig. 1b). Here, the simple RL models (Q1/Q2) only effectively learn about stimuli at the extremes of the ordered set (e.g., A and H, **supplementary Fig. S1a**), since these are statistically more likely to be winners or losers than their neighbours (under uniform sampling). No value learning occurs for intermediate items (stimuli B to G), since these are equally likely to be paired with lower and higher valued stimuli ^3^. However, the model can easily be adapted to performing transitive inference when extending it with a simple assumption: value updates should scale with the difference between the estimated item values, *Q*(A)-*Q*(B) (for similar approaches see refs. ^14,21,22^). More specifically, to the extent that A is already higher valued than B, observing the expected outcome A>B should induce weaker value updates, whereas the unexpected outcome A<B should induce stronger updates. To illustrate, observing an unknown amateur boxer win against a world champion should induce stronger changes in belief than the opposite, less surprising result (champion>amateur). When incorporating this simple assumption into our model (Model **Q1\***), it learns orderly structured values, *Q*(A) > *Q*(B) > … > *Q*(H), and thus accomplishes transitive inferences for all pairs of items (**Fig. 2d**; see also **supplementary Movie M1** for illustration how our Q-learning models accomplish transitive learning). We also observe a symbolic distance effect with this type of learning under partial feedback, similar to what we observed with simple RL under full feedback (cf. Fig. 2d and 2a).

Notably, the effect of asymmetric learning rates (*α*^+^ ≠ *α*^−^, Model **Q2\***) under partial feedback is strikingly different from what we observed with full feedback. Under partial feedback, optimal performance is achieved with a strongly asymmetric learning policy (α^+^>>α^-^ or α^+^<<α^-^), where only the winner (or loser) in a pair is updated (**Fig. 2e-f**, see also supplementary Movie M1). In other words, in a setting where hidden relational structure is inferred from only local comparisons, it is surprisingly beneficial to ignore losers (or winners) in outcome attribution. Of note, the winner/loser asymmetry outlined here differs from, and is orthogonal to, previously described asymmetries in learning from positive/negative ^e.g., 23–26^ or (dis-)confirmatory outcomes ^27,28^. A noteworthy aspect of our model Q2* is that the surprisingly superior, asymmetric learning policy results in a compression of the observer’s latent value structure (Fig. 2f). Selective updating therefore naturally gives rise to diminishing sensitivity towards larger values as is universally observed in Psychophysics ^29^, numerical cognition ^30,31^, and Behavioural Economics ^32^.

Going beyond typical studies of transitive inference with deterministic outcomes, we examined whether our simulation results generalize to scenarios where relational outcomes can be variable, as is the case in many real-world domains such as sports, stock markets, or social hierarchies. To this end, we added random variance to the comparison outcomes such that e.g., an item won over its lower-valued neighbour in approx. 80% of cases but lost in the other 20% (see *Methods* for details). Intuitively, we allowed for the possibility that competitor A may sometimes lose against B, even if A is generally stronger. We found that our simulation results held for such probabilistic environments, just as they did for deterministic scenarios (**supplementary Fig. S2)**.

In the models discussed so far, the observer associates each individual item (A, B, C, etc.) with an implicit value (*Q*(*A*), *Q*(*B*), *Q*(*C*), …; *item-level learning*). An alternative strategy is to more directly learn response preferences for each individual item *pairing* (*p*_*A*>*B*_, *p*_*B*>*C*_, …; *pair-level learning*; see *Methods*). For instance, in our partial feedback setting (Fig. 1), observers might learn to choose A when comparing A and B, to choose B when comparing B and C, and so forth, even without relying on value estimates for the individual items. In its simplest form, such memory for pairwise preferences (Model **P**) only allows learning of pair relations that had been directly experienced (i.e., only neighbouring pairs in our partial feedback setting; **supplementary Fig. S1b, *left***). However, the pair-level memory can also be extended to allow for transitive inference of more distant, never experienced relations ^8,33,34^ : when asked to judge e.g., A-C, observers might “chain together” memories of the linking neighbour preferences (*p*_*A*>*B*_ and *p*_*B*>*C*_) through associative recall ^17,35^ or spreading activation ^36^, to infer a transitive preference (*p*_*A*>*C*_; Model **Pi**; **supplementary Fig. S1b, *right***). Transitive inference based on such pair-level learning gives rise to an *inverse* symbolic distance effect (supplementary Fig. S1b, *right*), where nearby pairs are more discriminable than more distant pairs, reflecting the high dimensionality of the underlying associative memory structure. In modelling our empirical data, we allow for both item-level value learning (models denoted by a Q), pair-level learning (models denoted by a P), and a combination of both, in explaining human transitive inference.

## Results - Experiments

We report the results of four experiments (n=145) where we varied whether feedback was full or partial and whether it was probabilistic or deterministic (see *Methods*). In all experiments, participants were shown a pair of items (drawn from a set of 8) on each trial and were asked to make a relational choice (Fig. 1). Participants were given no prior knowledge about item values and could only learn through trial and error feedback.

### Full Feedback

In Experiment 1 (n=17), probabilistic choice feedback (see *Methods*) was provided after each of 448 sequential pair comparisons (“full feedback”). **Figure 3a** shows the mean proportions of correctly choosing the higher-valued item, averaged over all trials in Exp. 1. Descriptively, the choice matrix is dominated by a symbolic distance effect, as predicted by implicit value learning. Fitting our item-level learning models (Q1, Q2, Q1*, Q2*), the best fit of the data is provided by the simplest model (**Q1**) with a single learning rate for winners and losers (**Fig. 3c** and **3e**; protected exceedance probability: pxp(Q1)>0.99; mean BIC=361.79 ± 24.68 s.e.m.). In other words, participant behaviour was consistent with a symmetrical updating policy, which our simulations showed to be optimal in the full feedback setting.

**Fig. 3.**
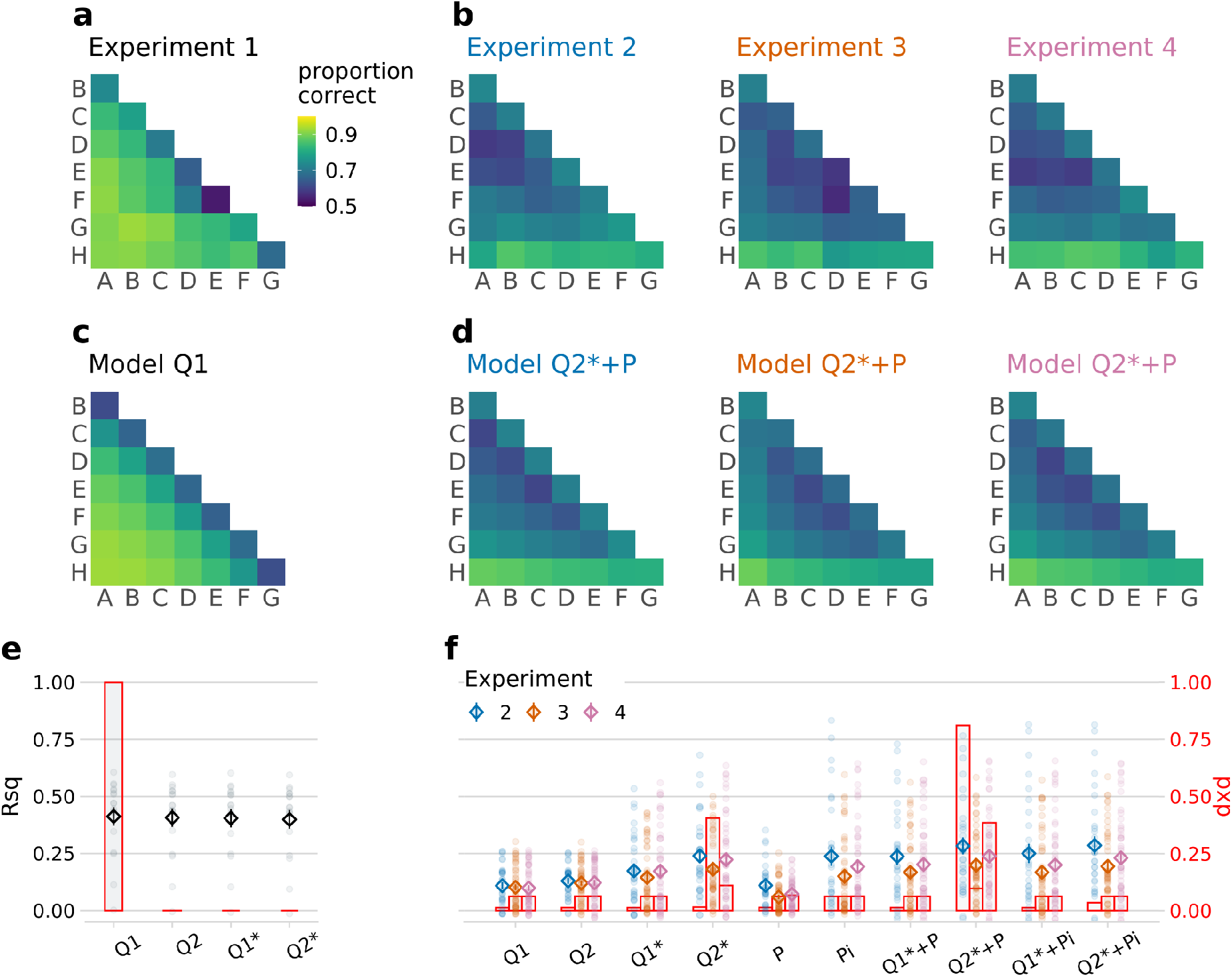
Empirical results and model fits. **a-b**, mean proportions of correct choices observed over all trials in each experiment (*a*, full feedback; *b*, partial feedback). **c-d**, mean choice probabilities predicted by the best fitting model in each experiment. **e-f**, model comparison (*e*, Exp.1; *f*, Exp. 2-4). Markers show model fits using a pseudo-R-squared (left y-axis, Rsq; diamonds and error bars show mean ± s.e.m., dots show individual participants). Rsq is inversely related to BIC, with larger values indicating better fit. Intuitively, Rsq=0 is equivalent to random chance, while Rsq=1 corresponds to a theoretically perfect model. Overlaid red bar graphs indicate each model’s probability of describing the majority of subjects best (right y-axis, pxp: protected exceedance probability, see *Methods*). The model space is described with the nomenclature **Q:** item-level learning; **1/2:** symmetric/asymmetric; **\*:** difference-weighted updating; **P**: pair-level learning; **i**: pair-relation-based inference.

### Partial Feedback

In Experiments 2-4, choice feedback was only provided on neighbour pairs (“partial feedback”) to study transitive inference. In these experiments, we increased the frequency participants were shown neighbouring pairs relative to non-neighbouring pairs to provide more learning opportunities, since the task is inherently harder. We verified that our simulation results were invariant to this modification (**supplementary Fig. S3)**. Otherwise, the design of Experiment 2 (n=31) was identical to Exp. 1. Experiment 3 (n=48) was an online replication of Exp. 2, where the pair items on each trial were shown side-by-side instead of sequentially. Experiment 4 (n=49) was similar to Exp. 3, but feedback was made deterministic (100% truthful) as in previous studies of transitive inference (see *Methods* for individual experiment details).

The choice data from each of the partial feedback experiments (Exp. 2-4, **Fig. 3b**) showed clear evidence for transitive inference, with above-chance performance for non-neighbouring pairs that never received feedback (mean accuracy averaged over non-neighbour trials, Exp. 2: 0.714 ± 0.028; Exp. 3: 0.698 ± 0.018; Exp 4: 0.709 ± 0.019; Wilcoxon signed-rank tests against chance level (0.5): all p<0.001, all r>0.84). Further, the grand mean choice matrices showed the following descriptive characteristics: (i) a symbolic distance effect similar to that observed with full feedback, (ii) an asymmetry with greater discriminability of lower-valued items, and (iii) relatively increased discriminability of neighbour pairs.

The modelling results for the partial feedback experiments are summarized in **Fig. 3d** and **3f**. We highlight two main findings. Firstly, the partial feedback data were better described by asymmetric models with different learning rates for winners and losers. This held true at every level of model complexity, with our asymmetric models (Q2, Q2*, Q2*+P, Q2*+Pi) always performing better than their symmetric counterparts (Q1, Q1*, Q1*+P, Q1*+Pi; Wilcoxon signed-rank tests comparing BICs, Exp. 2-4 combined: all p<0.001, all r>0.35), and regardless of whether the partial feedback was probabilistic (Exp. 2 and 3; comparison of mean BIC between asymmetric and symmetric models: both p<0.001, both r>0.60) or deterministic (Exp. 4; p<0.001, r=0.63). In other words, participants adopted an asymmetric learning policy, which proved superior in our model simulations (cf. Fig. 2e).

Secondly, behaviour in the partial feedback scenario was not fully described by item-level value learning alone. The winning model in Exp. 2 and 4 (Q2*+P; pxp=0.81 and 0.39; mean BIC=609.15 ± 12.77 and 434.27 ± 6.29) also incorporated pair-level learning, in addition to the value estimates of the individual items. This pair-level memory (+P, see *Methods*: *Models*) accounts for the increased performance for neighbouring pairs (Fig. 3b, first off-diagonals, cf. supplementary Fig. S1b). In Exp. 3, the model comparison was less clear, with model Q2* showing the highest pxp (0.41) but model Q2*+P providing a better average fit in terms of BIC (676.86 ± 19.40 vs. 692.09 ± 8.61, Wilcoxon signed-rank test: p<0.001, r=0.48). However, we found no evidence that pair-level memory contributed to transitive inference in our experiments. Incorporating associative recall of “linking” neighbour pairs (+Pi) worsened the model fits, both in terms of pxp (all pxp < 0.07) and BIC (Exp. 2-4 combined, Q2*+Pi: 570.42 ± 15.72 compared to Q2*+P: 567.60 ± 15.38; Wilcoxon signed-rank test: p<0.001, r=0.53), which is in line with the absence of an “inverse” symbolic distance effect (cf. Fig. S1b, *right*) in the empirical choice data (Fig. 3b).

Figure 4a illustrates how learning of non-neighbour comparisons in our experiments evolved over time. The value compression implied by asymmetric learning of winners (cf. Fig. 2e) predicts relatively better performance for lower-valued pairs (e.g., F-H) than for higher-valued pairs (e.g., A-C; **Fig. 4b**, *right*). We observed no such pattern in Experiment 1 with full feedback (Fig. 4a-c, *left*). In contrast, participants in Experiments 2-4 with partial feedback showed the critical pattern early on (Fig. 4a-c, *right*), as predicted by our asymmetric learning models (Fig. 4b, *right*). Turning to neighbouring pairs (**Fig. 4d**), which could additionally benefit from pair-level learning (+P, see above), our asymmetric model (Q2*+P) predicts only a modest decline of accuracy for higher-valued pairs (see also Fig. 3d), which also matched the empirical data (Fig. 4d, *right*).

**Fig. 4.**
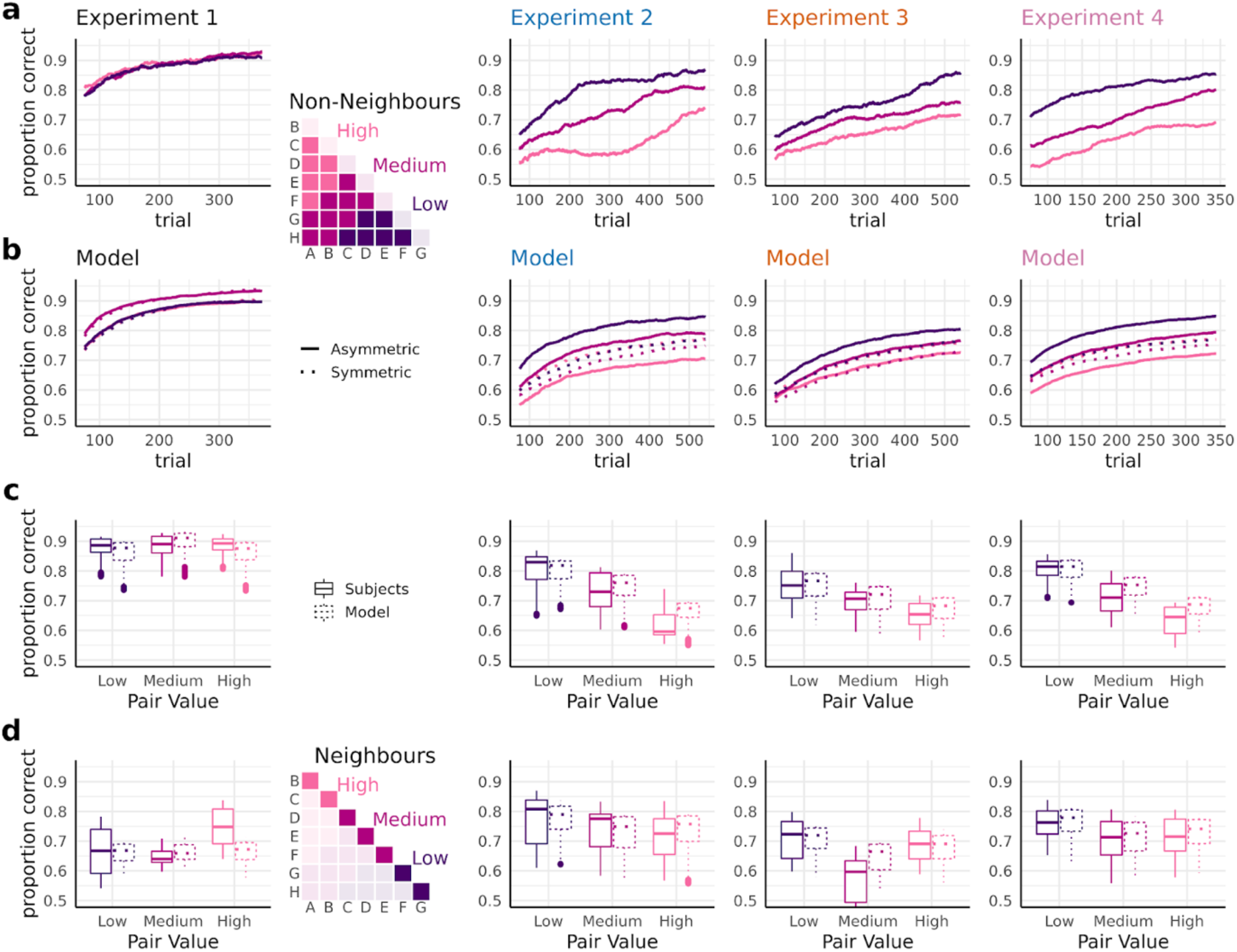
Learning curves and signatures of asymmetric learning. **a-b**, Learning of non-neighbour comparisons over time. **a**, Mean proportions of correct choices were calculated from a sliding window of 150 trials. Trajectories are shown separately for low-, medium-, and high-valued pairs (see inset matrix). **b**, Simulated learning curves using the best-fitting model in each experiment (see Fig. 3c-d). Solid lines: best-fitting asymmetric model; dashed lines: corresponding symmetric model with *α*^+^ = *α*^−^. Asymmetric learning in Exps. 2-4 is characterized by systematic performance differences (low > medium > high) as predicted by value compression (cf. Fig. 2f) early on in each experiment. **c**, Box plot summary of the results in *a*-*b*, averaged over all trials. **d**, same as *c*, for neighbouring item pairs.

The winner/loser asymmetries described so far might be explained by alternative learning biases, such as asymmetric learning weights for chosen vs. unchosen items ^37^. Given above-chance performance, the chosen item will statistically be more likely to be the winning item. To test this alternative explanation, we repeated our modelling analyses using separate learning rates (*α*^+^/*α*^−^) for chosen/unchosen items instead of for the winning/losing item (cf. *Methods*, equation 4). This alternative model fit our partial feedback data significantly worse (mean BIC collapsed across Exp. 2-4: 587.21 ± 15.91 vs. 567.60 ± 15.38, Wilcoxon signed-rank test p<0.001, r=0.54), corroborating our interpretation that transitive inference learning was better characterized by asymmetries between winners and losers. Previous RL-studies have also highlighted potential differences between learning from positive (confirmatory) as opposed to negative (disconfirmatory) feedback ^24,25,28^. Extending our winning model to incorporate such confirmation bias (see **Supplementary Fig. S5**) improved the overall model fit (mean BIC collapsed across Exp. 2-4: 537.91 ± 14.45 vs. 567.60 ± 15.38; Wilcoxon signed-rank test p<0.001, r=0.72), which is consistent with previous findings in other learning contexts ^24,25,28^. However, the addition of confirmation bias left the finding of winner/loser asymmetries unchanged (cf. Supplementary Fig. S5 and Fig. 3f), thus illustrating the robustness of our results.

We also compared our new model family against two previous models of transitive inference (see *supplementary Methods*): a classic value-transfer model (VAT) ^21^ and a more recent model based on ranking algorithms used in competitive sports such as chess (RL-ELO) ^22^. Both VAT and RL-ELO were outperformed by our winning model Q2*+P when fitted to our partial feedback data (Exp. 2-4 combined; mean BIC VAT: 606.59 ± 14.04; RL-ELO: 617.96 ± 15.07; Q2*+P: 567.60 ± 15.38; Wilcoxon signed-rank tests vs. Q2*+P: both p<0.001, both r>0.65) This held true even when we modified VAT and RL-ELO to include pair-level learning (+P) and separate learning rates for winners and losers (mean BIC=572.95 ± 15.17 and 576.34 ± 15.94, respectively; both p<0.014, both r>0.21). Thus, our asymmetric Q-learning process explains the experimental data better than these earlier models of transitive inference.

Our model simulations (Fig. 2e) indicate two aspects of asymmetric learning that are not directly evident from the group-level results shown in Figs. 3-4. First, performance benefits under partial feedback emerged not only for selective updating of winners, but likewise for selective updating of losers. Second, performance was highest for extreme asymmetries where the loser (or winner) in a pair was entirely ignored. We examined these aspects more closely on the individual participant level (**Fig. 5**). Half of the subjects in Exp. 2-4 (n=64) were indeed characterized by extreme asymmetry towards winners (with *α*^−^near zero). However, another subgroup (n=15) showed the opposite, an extreme asymmetry towards losers (with *α*^+^ near zero). In other words, in the partial feedback setting, most individuals showed an extreme bias towards winners or losers, either of which proved to be optimal policies in our model simulations (Fig. 2e and Fig. S2, *right*). In contrast, we found less substantial asymmetries under full feedback (Exp. 1) when allowing the learning rates for winners and losers to vary freely (i.e., using model Q2 instead of the winning model Q1). Statistical analysis confirmed that the asymmetries under full feedback (Exp. 1) were significantly lower than under partial feedback (Mann–Whitney U test of absolute asymmetry indices collapsed over Exp. 2-4 compared to Exp. 1: p=0.007, r=0.22; see *Methods*).

**Fig. 5.**
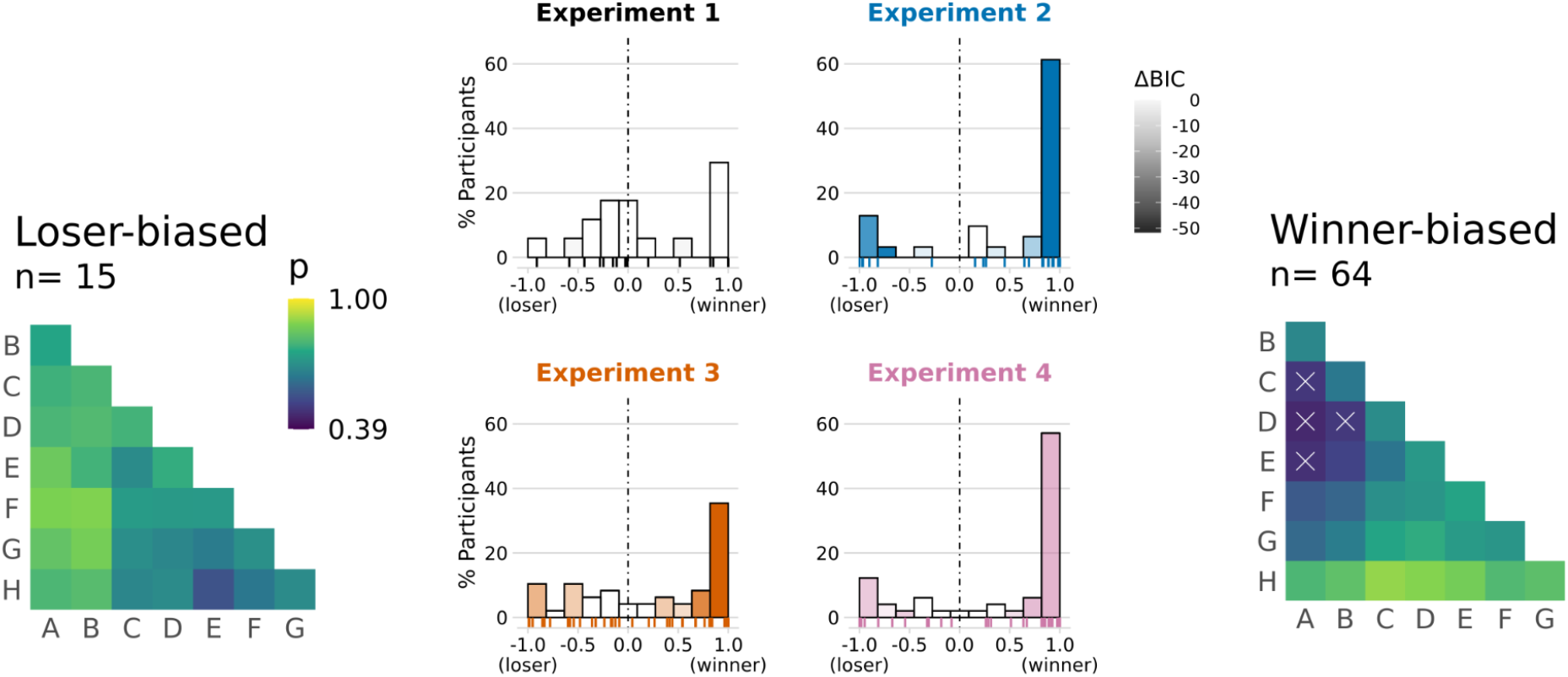
Winner/loser asymmetries in individual participants. *Middle*, histograms of participants in each experiment sorted according to normalized model-estimated asymmetry: (*α*^+^ − *α*^−^)/|(*α*^+^ *e α*^−^)|. Saturation of bars indicates improvement in model fit (*Δ BIC*, darker colours indicate greater improvement) compared to the corresponding symmetric model (i.e., Q1 in Exp. 1; Q1*+P in Exp. 2-4). No improvements can be seen in Exp. 1, where symmetric model Q1 provided the best fit (see also Fig. 3e). Raster plots on the bottom of each panel show individual participant results. The majority of participants (n=79 out of 128) in the partial feedback experiments (Exp. 2-4) showed strongly asymmetric updating either of winning or of losing items, with clear improvements in model fit. *Left*, mean choice behaviour of participants that were strongly biased towards losers (leftmost bars in *middle*, Exp. 2-4). p: proportion of correct choices. *Right*, same as *left*, for participants strongly biased towards winners (rightmost bars in *middle*). White crosses (x) indicate choice accuracies below chance (<0.50).

A potentially surprising observation in the subgroup of participants in Exp. 2-4 who selectively updated winners (Fig. 5 *right*) is a tendency for *below*-chance performance for relatively high-valued non-neighbours (e.g., A-D) despite each individual performing robustly above chance overall (cf. *Methods*: *Participants*). A potential explanation is that participants may sometimes have confused the two pair items in working memory at the time of feedback (cf. Fig. 1b, *right*). Such memory confusions would result in the items occasionally being updated with the incorrect learning rate and with incorrect sign (cf. *Methods*, equation 4). Under a learning policy that ignores losers, the losing items would then only be updated in error (and always incorrectly), resulting in a *negative* net learning rate for losers. Indeed, repeating our analysis while allowing for negative values of *α*^+^ and/or *α*^−^ yielded a small but significant improvement in model fit (mean BIC=562.52 ± 15.23 compared to 567.60 ± 15.38; Wilcoxon signed-rank test: p<.001, r=0.50). More specifically, in the *n*=64 participants who selectively updated winners (i.e., with a positive *α*^+^; mean=0.063 ± 0.007), *α*^−^ estimates were weakly negative (mean=-0.009 ± 0.0016; Wilcoxon signed-rank test against zero: p<0.001, r=0.64). Memory confusions may thus explain the systematically false inferences about certain item pairs (Fig. 5, *right*). Together, these findings are consistent with a strongly asymmetric learning mechanism that is also prone to occasional memory errors.

To summarize our empirical findings, when transitive relations could only be inferred from local comparisons (Exp. 2-4), human learning was characterized by an asymmetric outcome attribution to either winners or losers, which proved to be surprisingly optimal in model simulations. In contrast, a symmetric attribution of relational outcomes emerged in a setting where all pair relations could be directly experienced (Exp. 1), and for which our simulations identified symmetric updating to be the most efficient.

## Discussion

Reasoning about the relationships between arbitrary pairings of items is a key component of human intelligence. Through simulations, we showed how different learning regimes perform better in full and partial feedback contexts. Under full feedback, the best learning model used *symmetric* learning to update the value estimates for the winning and losing items in opposite directions, with the same magnitude. However, under partial feedback (only for neighbouring items), the best learning model used *asymmetric* learning to only update the value representations for either the winner or the loser. Across four experiments, we find robust evidence that human learners used the best learning rule to match their feedback context. Participants used symmetric learning under full feedback (Exp. 1) and asymmetric learning under partial feedback (Exp. 2-4). While our asymmetric models allowed for a wide range of possible learning rate combinations, a majority of subjects showed one-sided learning, where value representations were only updated for either winners or losers.

An important feature found both in our model simulations and participant behaviour is a compression of the emerging implicit value structure, which results in a systematic decrease in discriminability of higher valued items (see Fig. 2f and Fig. 4). This resembles the Weber-Fechner Law in psychophysics ^29^, where sensitivity to stimulus differences diminishes with increasing magnitude (see also refs. ^38,32^). While there exist alternative theoretical accounts for this ubiquitous phenomenon ^39^, our findings add a new perspective: compressed representations of magnitude emerge naturally from a learning policy that is optimized for inferring global relationships from local comparisons. From this perspective, subjective compression might not only reflect an efficient adaptation to the distribution of stimuli in the environment ^40–42^, but could also result from learning policies that enhance transfer to novel relationships.

In other contexts, previous RL studies have discovered different types of learning asymmetries, such as between positive and negative ^24,25^ or confirmatory and disconfirmatory outcomes ^28^. The one-sided learning policy highlighted here in the context of transitive inference is orthogonal to these other asymmetries, but may play a similar role in leveraging a biased but advantageous learning strategy (see also refs. ^27,43,44^). Unlike with *optimal* cognitive biases reported previously ^45–49^, we did not find the benefit of the present learning asymmetries to emerge from general limitations (noise) in decision making (**supplementary Fig. S4**). We speculate that human learners may adopt the present biases more strategically, in settings where the availability of only sparse feedback presages the requirement of future inferential judgments.

Previous theories have proposed richer and more complex cognitive mechanisms for transitive inference, often with an emphasis on the key role of the hippocampus in representing relational knowledge ^15,50^. Early research appealed to the idea that individuals used spatial representations to learn ordered value sequences ^1,8,51^. More recently, various models have been proposed that use associative learning mechanisms to describe how interactions between episodic memories in the hippocampus can generalize relational knowledge from local to distant comparisons ^17,52^. In our present experiments, we found no evidence for transitive inference through such “associative linking” and failed to observe its key empirical prediction (an inverse symbolic distance effect, cf. Fig. 3b and S1b, *right*). We show instead that simpler mechanisms of value learning ^21,53,54^ combined with clever biases (i.e., asymmetric learning rates) can be sufficient for performing TI and for accurately describing human learners.

In summary, we report evidence for pronounced asymmetries in transitive relational learning, where observers selectively update their beliefs only about the winner (or the loser) in a pair. Although asymmetric learning yields distorted value representations, it proves beneficial for generalization to new, more distant relationships. Thus, this biased learning regime appears well-adapted for navigating environments with relational structure on the basis of only sparse and local feedback.

## Methods

### Participants

Participants in Exp. 1 and 2 were recruited from a participant pool at the Max Planck Institute for Human Development. Of these, n=20 participated in Experiment 1, (13 female, mean age 27.15 ± 3.91 years) and n=35 participated in Experiment 2 (14 female, 27 ± 3.80 years). Participants in Exp. 3 and 4 were recruited online via Prolific Academic (www.prolific.co) with n=76 completing Exp. 3 (23 female, 24.73 ± 5.40 years) and n=60 completing Exp. 4 (23 female; 25.92 ± 4.54 years). Participants in Exp. 1 and 2 received a compensation of €10 per hour and a bonus of €5 depending on performance. Payment in Experiment 3 and 4 was £4.87 (£1.46 bonus) and £3.75 (£1.12 bonus), respectively. We obtained written informed consent from all participants and all experiments were approved by the ethics committee of the Max Planck Institute for Human Development.

Participants who did not reach above-chance learning levels were excluded from analysis. The threshold for inclusion was set to 60% correct judgments in the last two blocks of the experiment, which corresponds to a binomial test probability of p<0.01 compared to chance-level (50%). After exclusion, n=17 (Exp. 1), n=31 (Exp. 2), n=48 (Exp., 3) and n=49 (Exp. 4) participants remained for analysis.

### Stimuli, task, and procedure

In Exp. 1 and 2, eight pictures of everyday objects and common animals were used as stimuli (Fig. 1a). In Exp. 3 and 4, we included 12 additional pictures of objects and animals and selected for each participant a new subset of 8 images as stimuli. An additional set of 8 pictures was used for instructions and practice purposes in each experiment. All images were from the BOSS database ^55^, with the original white background removed.

All experiments involved learning the latent relations between the 8 stimuli (A>B>C>D>E>F>G>H) through pairwise choice feedback, where the latent value structure was pseudo-randomly assigned to the pictures for each participant. On each trial, a pair of pictures was presented and observers were asked to choose the higher-valued stimulus (two-alternative choice with time-out). All possible stimulus pairings (8 neighbours and 20 non-neighbours) were randomly intermixed across trials, with randomized ordering of the elements in a pair (e.g., A-B or B-A). Prior to all experiments, participants were given written instructions and were asked to complete two brief practice blocks to familiarize with the task.

### Experiment 1 (full Feedback, n=17)

On each trial, two items were presented one after the other at fixation (0.5 s/item) with an inter-stimulus interval of 2-3s (randomized). After the 0073econd item, Arabic digits “1” and “2” were displayed to the left and right of fixation (positions randomized across trials) and participants were asked to choose the higher-valued item by pressing the corresponding arrow key (left or right) within 2s. A written feedback message (“great” for correct responses, “incorrect” for errors) was shown after each choice (neighbouring and non-neighbouring pairs). The items’ latent values in Exp. 1 were probabilistic (with a Gaussian distribution) and designed such that feedback was truthful on approx. 80% of neighbour trials (“probabilistic feedback”). Each Participant performed 448 learning trials with all possible stimulus pairings (n=56) presented in each of 8 consecutive blocks. Experiments 1 and 2 were conducted in lab, using Psychophysics Toolbox Version 3 ^56^ running in MATLAB 2017a (MathWorks).

### Experiment 2 (partial feedback, n=31)

The design was nearly identical to Exp. 1, but choice feedback was only given after neighbouring pairs. After non-neighbouring pairs instead, a neutral “thank you” message was displayed. Neighbouring pairs were presented more often (2.5 times as often as non-neighbouring pairs), resulting in 616 trials (presented in 8 blocks of 77). In Exp. 2, we additionally recorded EEG and participants performed a brief picture viewing task prior to the experiment. These data were collected for the purpose of a different research question and are not reported here.

### Experiment 3 (partial feedback, n=48)

The basic design was identical to Exp. 2, except for the following changes: Both pair items were displayed simultaneously on screen for 2.5 s, one to the left and the other to the right of a centred fixation cross. Participants were instructed to quickly select the higher valued item using the left or right arrow key. After neighbouring pairs, a feedback message (“win” or “loss”) was presented. After non-neighbouring pairs, no feedback message was shown. Experiments 3 and 4 were programmed in PsychoPy 2020.1.3 ^57^ and conducted online (Pavlovia.org), with intermittent attention checks.

### Experiment 4 (partial feedback, deterministic, n=49)

The design was identical to Exp. 3, but feedback was always truthful (“deterministic feedback”). As learning expectedly proceeds faster with deterministic feedback, neighbouring pairs were presented only 2 times as often as non-neighbours and we reduced the number of trials to 420 (presented in 6 blocks of 70 trials).

## Models

### Item-level learning

To model how observers update their value estimates about the winning item *i* and the losing item *j* after relational feedback, we assume a simple delta rule ^58^ (Model **Q1**):

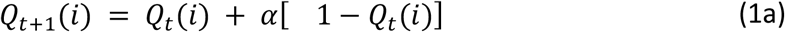

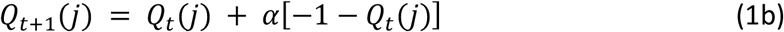

where *Q*_*t*_ is the estimated item value at time *t* and *α* is the learning rate.

Transitive inference is enabled by a modified updating rule ^similar to 22,14^ based on the relative difference *d*_*t*_(*i, j*) between the value estimates for the winner *i* and the loser *j* in a pair:

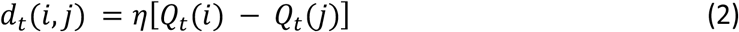

where *η* is a scaling factor. Value updating is then moderated by the extent to which feedback is consistent (or inconsistent) with *d*_*t*_(*i, j*) (Model **Q1\***):

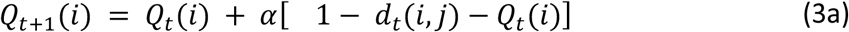

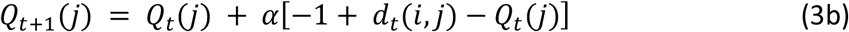

for the winning item *i* and the losing item *j*, respectively. Note that equation 1 is a special case of equation 3 when *η* = 0.

We can allow asymmetric updating of winners and losers by introducing separate learning rates *α*^+^ and *α*^−^ (Models **Q2**/**Q2\***):

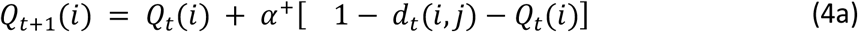

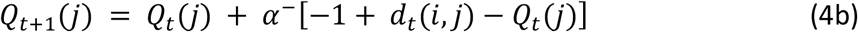

where the winning item *i* is updated via *α*^+^ and the losing item *j* is updated via *α*^−^.

To convert the value estimates from item-level learning into pairwise choice probabilities for any two items *i* and *j*, we use a logistic choice function to define the probability of choosing *i* > *j* based on the difference between the estimated item values:

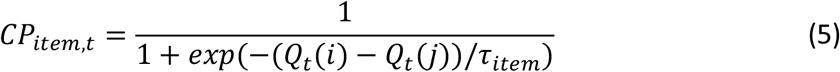

where *τ*_*item*_is the (inverse) temperature parameter controlling the level of decision noise in choices based on item-level learning.

### Pair-level learning

For the partial feedback scenario, we also define an alternative learning model (Model **P)** that learns pairwise preferences between neighbouring items (rather than the individual items’ values). For each neighbouring pair *n* (1..7), we can describe the preference between its members (e.g., *p*_*A*>*B*_) probabilistically in terms of a beta distribution:

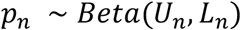

Following truthful feedback (e.g., “correct” when A>B was chosen), the upper value of the beta distribution is updated, increasing the preference in favour of the higher-ranking pair member:

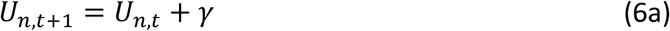

whereas following untruthful feedback (only in experiments with probabilistic feedback, see Exp. 2 and 3), the lower value is updated, reducing the preference for the higher-ranking member:

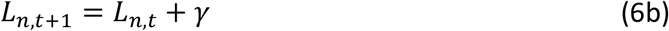

with *γ* acting as a learning rate. We can thus define the learned neighbour preference at time *t* based on the expectation of the beta distribution (Model **P**):

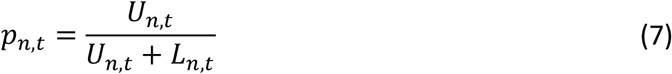

where *p*_*n,t*_ = 0.5 reflects indifference and values of *p*_*n,t*_ larger (or smaller) than 0.5 reflect a preference for the higher (or lower) ranking pair member. While this mechanism can learn the relations between neighbouring items under partial feedback, it fails to learn the relations between non-neighbouring items for which there is no direct feedback signal. However, transitive inference of preferences between non-neighbouring items is possible through associative recall of those neighbour preferences that “link” the two non-neighbour items in question. To allow for this possibility, we define the inferred preference between any two items *i* and *j* via the intermediate neighbour preferences *p*_*n,t*_ ∈ *M* separating *i* and *j* (Model **Pi**):

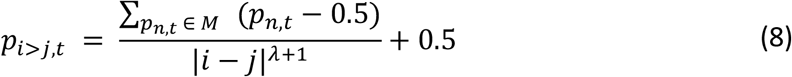

where |*i* − *j*| is the rank distance between the items’ true values, and *λ* is a free parameter reflecting failure to retrieve linking pair preferences in the range [0, ∞]. If *λ* = 0, preferences between non-neighbours will be a lossless average of all intermediate neighbour preferences (i.e., perfect memory). As *λ* grows, the preference between non-neighbours will shrink to indifference with increasing distance between *j* and *i*. In other words, this model performs perfect transitive inference if *λ* = 0, and no transitive inference as *λ* → ∞. Note that for neighbour pairs (where |*i* − *j*| = 1), equation 8 is equivalent to equation 7.

We again use a logistic choice rule to define the probability of choosing item *i* over *j* based on pair preference *p*_*i*<*j,t*_ subject to decision noise *τ*_*pair*_:

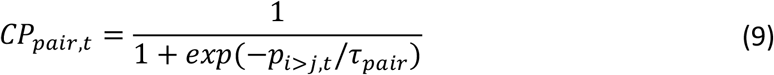

From equations 6-9, we constructed alternative models incorporating basic pair-level learning (Model **P**, where *λ* is fixed at a large value) and pair-level transitive inference (Model **Pi**, where *λ* is a free parameter).

To combine item-level (equations 1-5) and pair-level (equations 6-9) learning, we assume that choices are triggered by whichever of the two models provides a stronger preference on a given trial. Thus, choices are based on item-level learning (*CP*_*item*_) if:

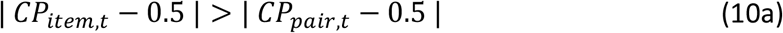

and are based on pair-level learning (*CP*_*pair*_) if:

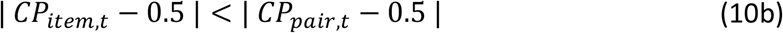

This effectively implements a mixture of item- and pair-level learning.

### Model space

From equations 1-10, we constructed a nested model space (see Supplementary Fig. S1c) with either one or two learning rates (**1**: symmetric, **2**: asymmetric updating, cf. equation 4). One set of models allows for simple item-level RL only (Models **Q1** and **Q2**) or additionally for item-level transitive inference (Models **Q1\*** and **Q2\***, equations 1-5). Alternative models (equations 6-9) incorporated pair-level learning (Model **P**) and pair-level inference (Model **Pi**). Mixture models (equation 10) combined item-level and pair-level learning, specifically (**Q1*+P, Q2*+P, Q1*+Pi, Q2*+Pi**). Technically, all models under study were derived from the most flexible model Q2*+Pi with individual parameter restrictions (e.g., *γ* = 0 yields model Q2*; or *α*^+^ = *α*^−^ yields symmetric updating).

### Performance simulations

We simulated the performance of our item-level learning models (Q1, Q2, Q1*, Q2*) in tasks akin to those used in the human experiments, with full and partial feedback (Fig. 2 and Fig. S1-S2). The performance simulations were run in Matlab R2020a (MathWorks). Models were initialized with flat priors about the item values (all *Q*_1_(*i*) = 0, i.e., the first choice was always a random guess with *CP*_1_ = 0.5). Like in the human experiments, choice feedback was provided either for all pairs (full feedback) or only for neighbour pairs (partial feedback). We simulated model performance over a range of learning rates (*α*^+^ and *α*^−^, 0 to 0.1 in increments of 0.001). Relational difference-weighting (*η*) was set to either 0 (models Q1/Q2) or 8 (models Q1*/Q2*), and decision noise (*τ*_*item*_) was set to 0.2 and 0.04 (full and partial feedback) which resembles the noise levels estimated in our human observers in the respective experiments. Mean choice probabilities (e.g., Fig. 2a *lower*) and performance levels (e.g., Fig 2b) were simulated using the same number of trials and replications (with a new trial sequence) as in the respective human experiments. Simulation results under partial feedback (Fig. 2e and S2-4) were qualitatively identical when inspecting performance on non-neighbouring pairs only.

### Parameter estimation and model comparison

Model parameters were estimated by minimizing the negative log-likelihood of the model given each observer’s single-trial responses (from all trials in the experiment) across values of the model’s free parameters [within bounds (lower;upper): *α*/*α*^+^/*α*^−^(0;0.2), *η*(0;10), *τ*_*item*_ (0;1), *γ*(0;1), *λ*(0;100), *τ*_*pair*_(0;1), with a uniform prior]. The best-fitting parameter estimates are shown in **supplementary Fig. S6**. Model fitting was performed in R (R core team, 2020; https://www.R-project.org/). Minimization was performed using a differential evolution algorithm ^59^ with 200 iterations. We then computed the Bayesian Information Criterion (BIC) of each model for each participant and evaluated the models’ probability of describing the majority of participants best (protected exceedance probability, pxp) ^60^. In Fig. 3e and 3f, we also provide a Pseudo-R-squared computed as *Rsq* = 1 − (*BIC* _,*model*_ *BIC* _*null*_), which quantifies goodness of fit relative to a null model of the data, with larger values indicating better fit (similar to ref ^61^). Model comparisons for Exp.1 (full feedback) were restricted to item-level learning models, as the availability of direct feedback for every pairing would equate pair-level learning models (P, Pi) to homogenous learning of all pairs, obviating contributions from transitive inferences.

To quantify model-estimated asymmetry (Fig. 5), we computed an index of the normalized difference in learning rates *A* = (*α*^+^ − *α*^−^)*/*|(*α*^+^ + *α*^−^)| which ranges from -1 (updating of losers only) to 1 (updating of winners only), with *A* = 0 indicating symmetric updating. For comparison between full and partial feedback experiments, we contrasted the absolute |*A*| estimated from the winning model in Exp. 2-4 (Q2*+P, see Fig. 3f) with that estimated from model Q2 in Exp. 1.

### Model- and parameter recovery

To establish whether the individual models can be distinguished in model comparison we simulated, for each participant and model, 100 experiment runs using the individuals’ empirical parameter estimates under the respective model. We then fitted the generated data sets (binomial choice data) with each model and evaluated how often it provided the best fit (in terms of BIC). This way we estimated the conditional probability that a model fits best given the true generative model [*p*(*fit*|*gen*)]. However, a metric more critical for evaluating our empirical results is *p*(*gen*|*fit*), which is the probability that the data was generated by a specific model, given that the model was observed as providing the best fit to the generated data ^62^. We compute this probability using Bayes theorem, with a uniform prior over models [*p*(*gen*)]:

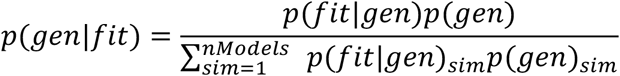

To mimic the level of inference in our human data fitting, we examined mean *p*(*fit*|*gen*) and *p*(*gen*|*fit*) on the experiment level, based on full simulations of all participants in Exp. 1 (full feedback) and Exp. 2 (partial feedback). Critically, under partial feedback (cf. Exp. 2-4), all our models were robustly recovered with this approach (**supplementary Fig. S7**).

Under full feedback (Exp. 1), human participant behaviour was best characterized by symmetric learning rates (*α*^+^ ≈ *α*^−^), even when both learning rates were free parameters (Fig. 3e and Fig. 4). To test whether we could have detected asymmetric learning, had it occurred in Exp. 1, we enforced asymmetry in simulation by setting *α*^−^ to values near zero (by drawing from a rectified Gaussian with *µ* = 0 and *σ* = 0*n*01). We likewise enforced difference-weighted updating (*η* > 0) when simulating model Q2*, by setting *η* to similar levels as empirically observed in the partial feedback experiments (*µ* = 3 and *σ* = 0*n*5). With this, the model recovery for Exp. 1 successfully distinguished between symmetric (Q1/Q1*) and asymmetric learning models (Q2/Q2*, **supplementary Fig. S8**). However, models with difference-weighted updating (Q1*/Q2*, equations 2-3) were partly confused with models Q1/Q2. In other words, our empirical finding of Q1 as the winning model in Exp. 1 (Fig. 3e) does not rule out the possibility of Q1* as the generative process under full feedback.

To establish whether our inferences about model parameters (e.g., Fig. 5) are valid, we simulated choices under partial feedback (Exp. 2) using our winning model (Q2*+P). Choice data sets were simulated using each participant’s empirical parameter estimates and iteratively varying each parameter over 20 evenly spaced values within the boundaries used in *Parameter estimation* (see above). We then fit the model to the simulated data sets and examined the correlations between generative and recovered parameters (**supplementary Fig. S9 and S10**). All fitted parameters correlated most strongly with their generative counterparts (min 0.59, max 0.93) while correlations with other generative parameters were generally weaker (min -0.44, max 0.43).

### Statistical analyses

Behavioural and modelling results were analysed using nonparametric tests (two-sided) as detailed in *Results*. In case of multiple tests, the maximum p-value (uncorrected) is reported.

## Code and data availability

The data that support the findings of this study are available at: https://arc-git.mpib-berlin.mpg.de/ti/asymm

The experiment- and analysis code are available at: https://arc-git.mpib-berlin.mpg.de/ti/asymm

## Acknowledgements

We thank Stephanie Nelli, Christopher Summerfield, and Nico Schuck for helpful feedback and discussion, and Stefan Appelhoff and Jann Wäscher for technical support.

This work was supported by a DFG (German Research Foundation) research grant to BS (DFG SP-1510/6-1). CMW is supported by the German Federal Ministry of Education and Research (BMBF): Tübingen AI Center, FKZ: 01IS18039A and funded by the Deutsche Forschungsgemeinschaft (DFG, German Research Foundation) under Germany’s Excellence Strategy – EXC 2064/1 – 390727645.

## Author contributions

JLD and BS designed the experiments. JLD, CW, and IP performed the experiments. BS designed the modelling approach. SC and BS performed the simulations and analyses with contributions from CMW and JLD. SC and BS visualized the simulations and results. BS, CMW, SC, and JLD wrote the paper.

## Supplementary Movie

**Supplementary Movie M1.**
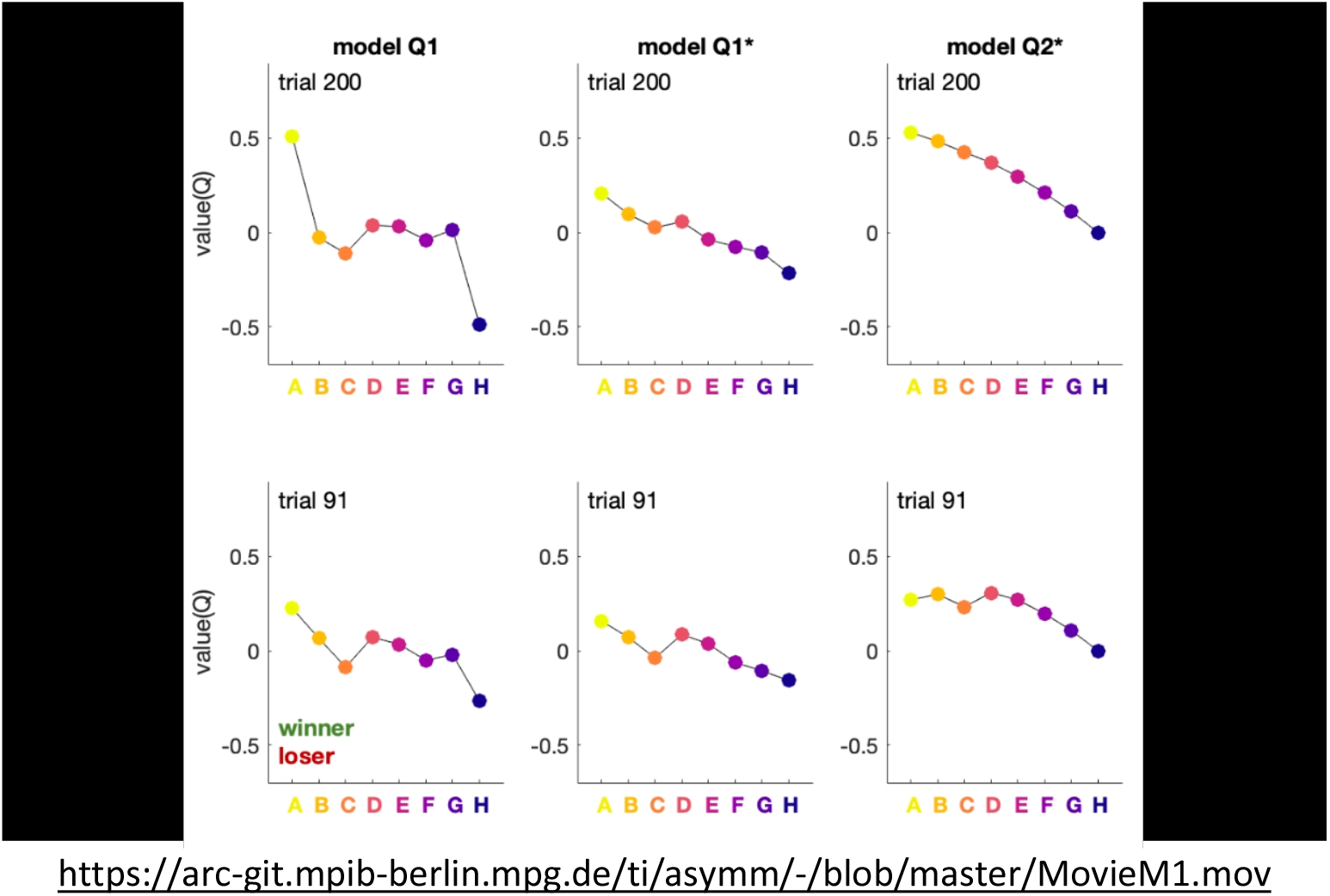
Simulations of Q-learning under partial feedback for models Q1 (*left*) and Q1* (*middle*) with the same learning rate for winning and losing items, and for model Q2* (*right*) with asymmetric learning about winners only (*α*^−^ set to 0). Colored dots indicate the momentary Q-values for items A-H. Simulations are shown for a trial sequence with deterministic feedback for illustration purposes. The movie first plays 200 learning trials (top panels) and then repeats the same trials more slowly (bottom panels). Non-neighbour trials (on which no feedback is given) are fast forward. Green and red disks indicate the winning and losing item on every trial. Model Q1 (*left*) effectively learns only about the extreme items (A and H), while intermediate item values fluctuate unsystematically around the pre-experiment baseline. In models with difference-weighted updating (Q1* and Q2*, *middle* and *right*), value differences propagate through the item series, which leads to a more monotonic value structure that enables transitive inferences also about intermediate items (B-G). In model Q1* (*middle*) with symmetric learning, propagation can occur in both directions, which results in partly conflicting updates for mid-range items (e.g., C-F). This induces residual non-monotonicity in the evolving value structure, which can compromise transitive inference. In model Q2* (*right*) with asymmetric learning, in contrast, conflicting updates are reduced, leading to a more strictly monotonic value structure that enables superior transitive inference.

## Supplementary Figures

**Supplementary Figure S1.**
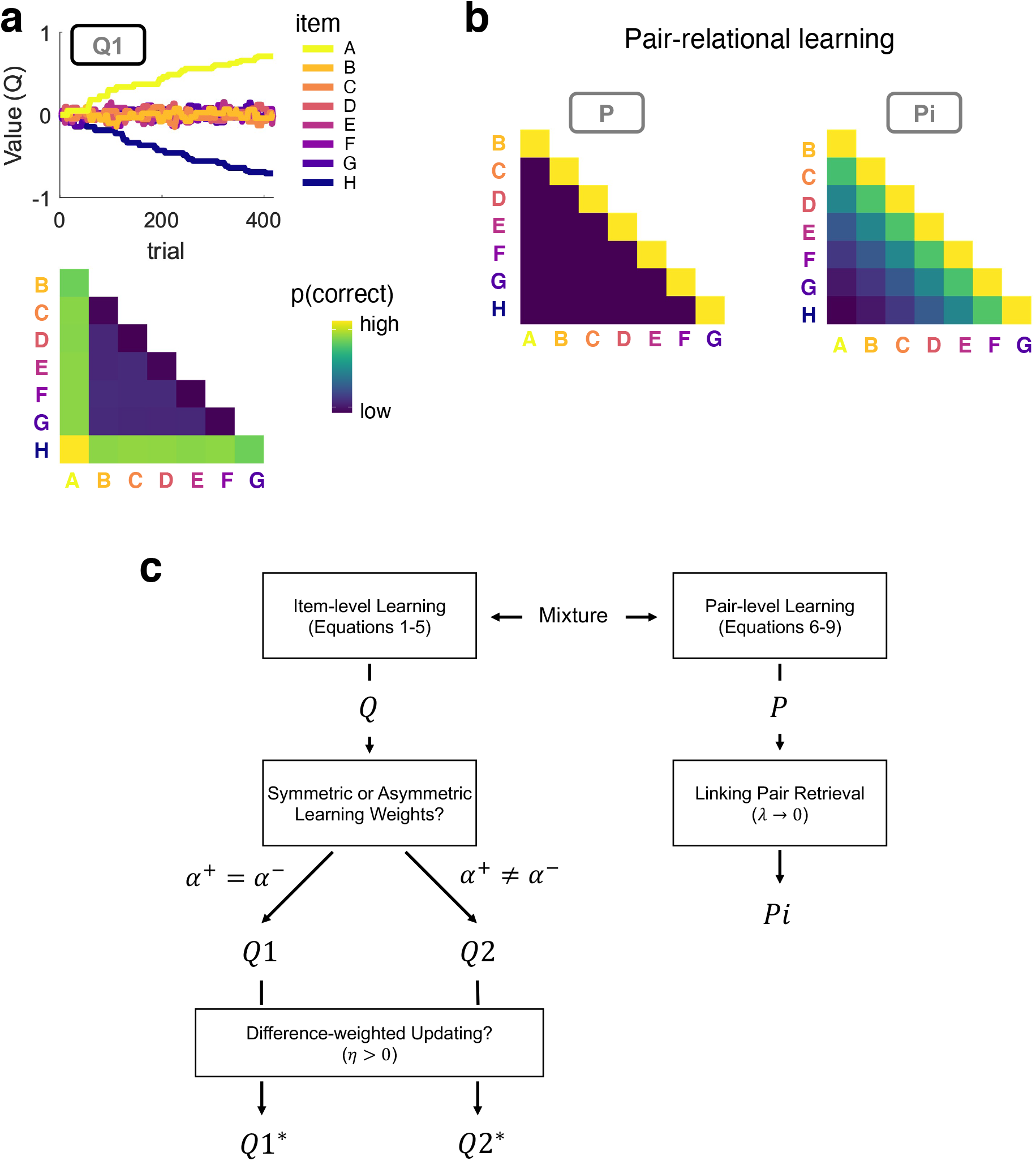
**a**, Simulation of model Q1 under partial feedback. Same conventions as in Fig. 2. The simple Q-learning models (Q1/Q2) can only learn about the extreme items (here, A and H) under partial feedback. **b**, Choice matrices predicted by pair-level learning without (*left*, model P) or with associative recall of “linking” pair relationships (*right*, model Pi). Choice behaviour was simulated with a pair-level learning rate *γ* = 1. Associative recall in model Pi (*right*) was enabled by additionally lowering parameter *λ* to 1 (see *Methods* for details). **c**, Schematic overview of the model space. See *Methods*, Models.

**Supplementary Figure S2.**
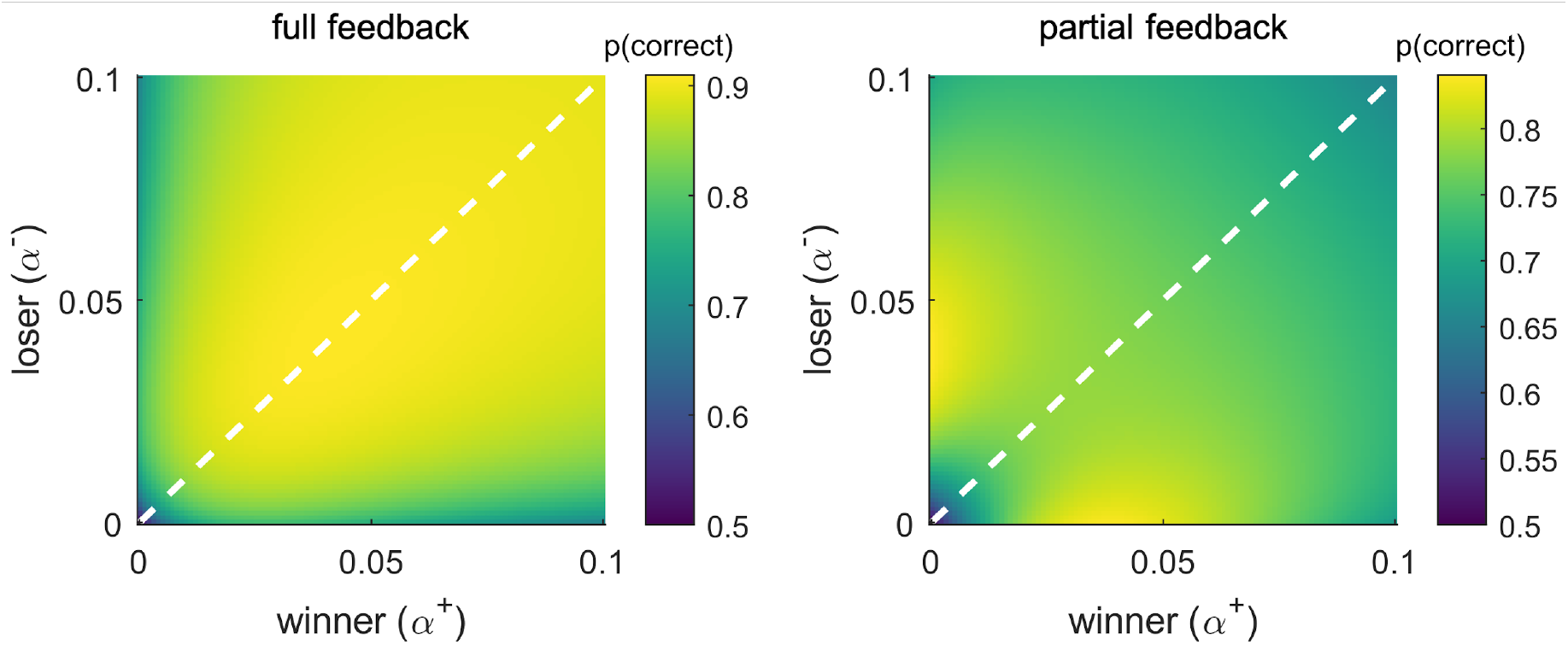
Performance simulations with probabilistic choice outcomes, same conventions as in Fig. 2. *Left*, full feedback (for all pairs; cf. Fig. 2b). *Right*, partial feedback (only for non-neighbouring pairs; cf. Fig. 2e). Optimal learning under partial feedback is characterized by asymmetric updating (*α*^+^ ≠ *α*^−^), just as was observed with deterministic outcomes (cf. Fig. 2e).

**Supplementary Figure S3.**
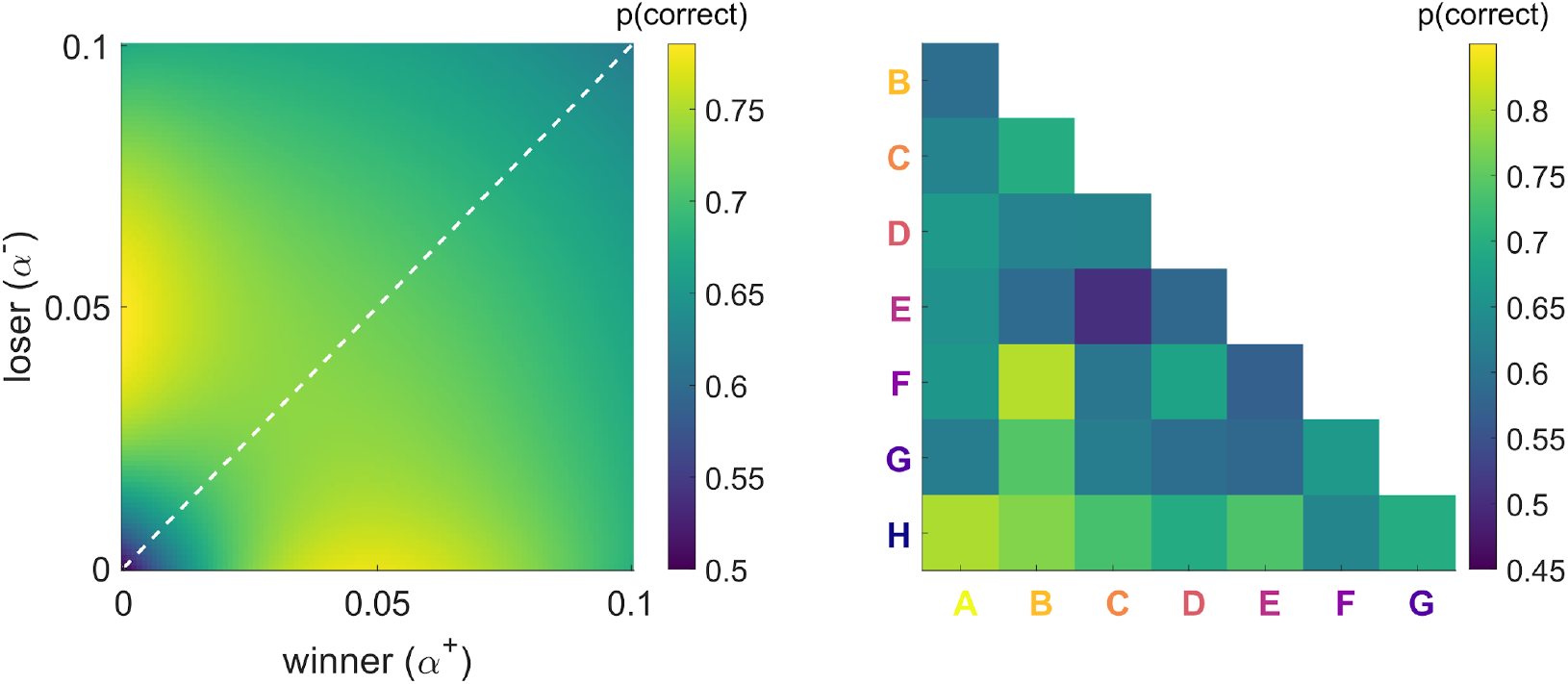
Pilot experiment with partial feedback (n=11). The design was identical to Exp. 2, except that all item pairs (neighbours and non-neighbours) were presented equally frequently (like in Exp. 1). *Left*, Performance simulation shows a similar benefit of asymmetric updating as we observed in simulation of Exp. 2-4 (where neighbouring pairs were presented more frequently, cf. Fig 2e and S2, right). *Right*, Mean proportions of correct choices in the pilot experiment. The overall learning level was relatively low, with n=9 (of 20) pilot participants not meeting our inclusion threshold for above-chance performance (cf. *Methods: Participants)*. The descriptive choice data of the remaining 11 pilot participants (shown in *right*) indicate a similar learning asymmetry as we observed in our main experiments.

**Supplementary Figure S4.**
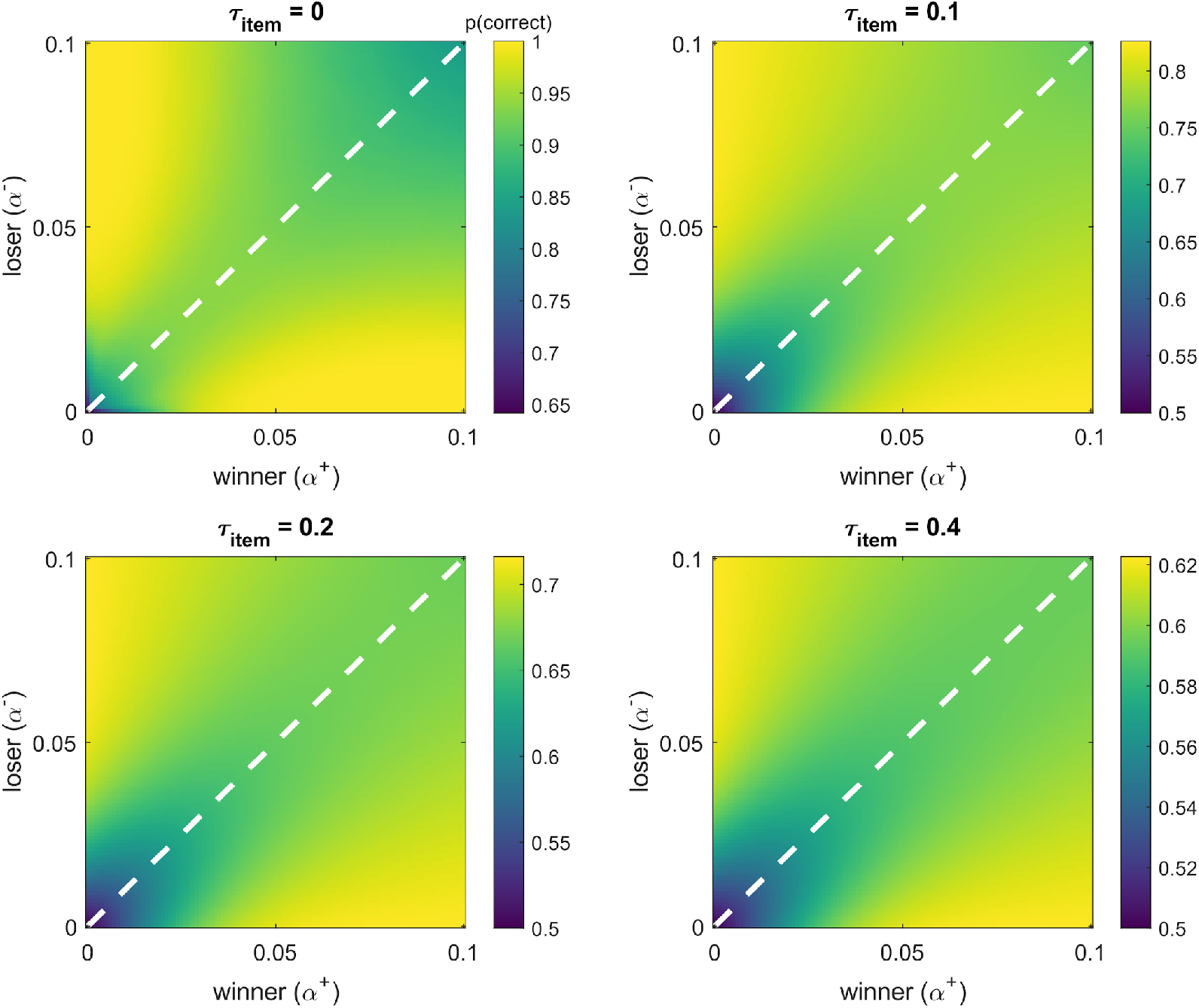
Performance simulations under partial feedback (analogous to Fig. 2e) for different levels of decision noise (*τ*_*item*_). Asymmetric learning is beneficial regardless of decision noise level and accordingly, across a wide range of overall performance levels. Note that different colour scales for each panel are used to increase interpretability (see colour bars). Simulations with probabilistic outcomes yielded a qualitatively very similar pattern.

**Supplementary Figure S5.**
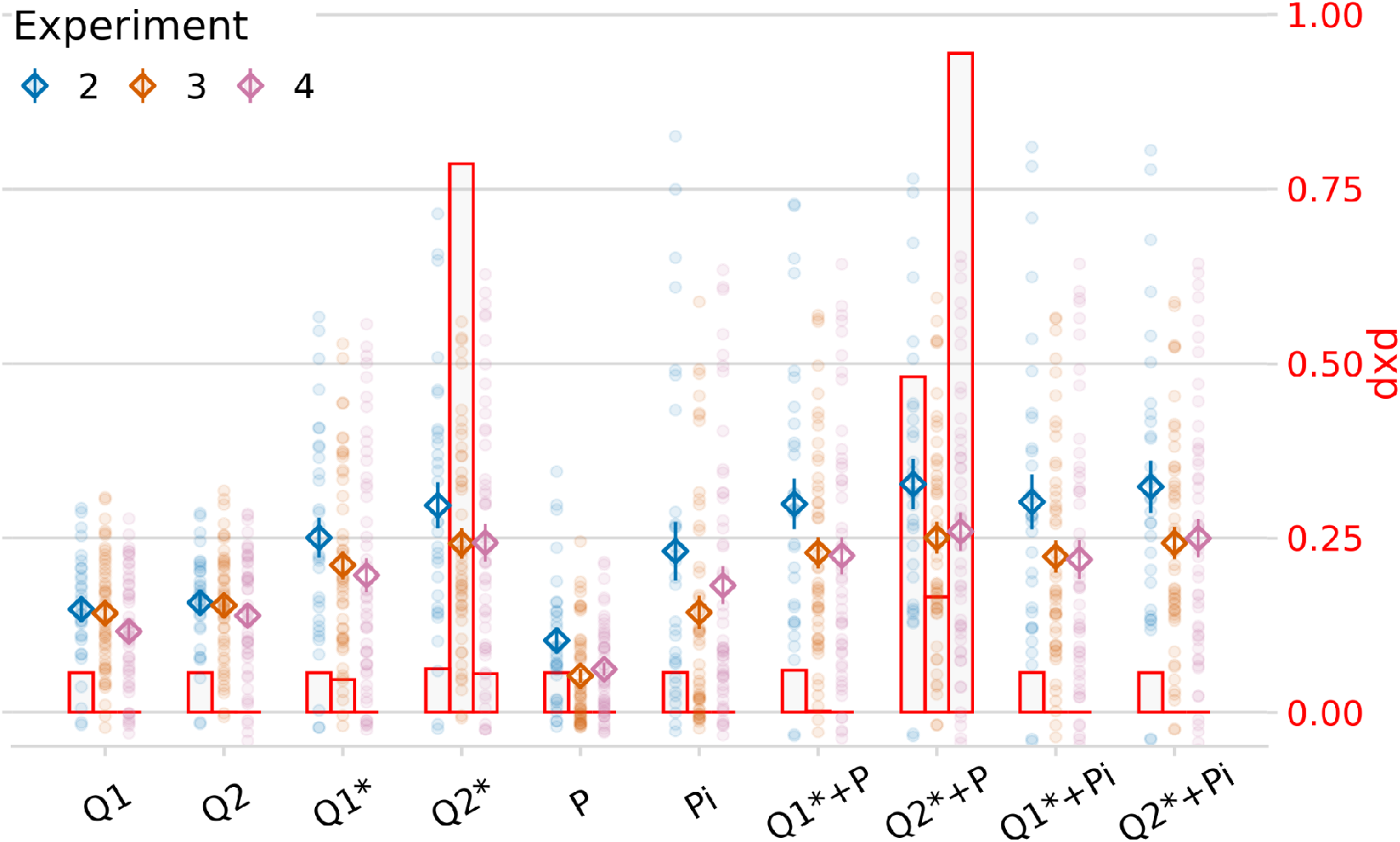
Model comparison (Exp. 2-4) analogous to Fig. 3f, but additionally allowing for differential learning from confirmatory vs. disconfirmatory choice feedback. Here, all models included an additional parameter *ω* ∈ (0;10), by which belief-confirming learning rates were modelled as: *α*_*conf*_ = *α* * *ω*. Same conventions as in Fig. 3f. Markers show model fits using a pseudo-R-squared (Rsq, left y-axis; diamonds and error bars show mean ± s.e.m., dots show individual participants). Overlaid red bar graphs indicate each model’s probability of describing the majority of subjects best (right y-axis, pxp: protected exceedance probability). While the extra parameter *ω* led to general improvements in fit (note overall higher Rsq compared to Fig. 3f), the model comparison result with respect to winner/loser asymmetries was identical, both in terms of Rsq/BIC and pxp. The estimates of parameter *ω* in the winning models were larger than 1 (mean=4.72, SE=0.27, p<0.001, r=0.83, Wilcoxon signed-rank test against 1, collapsed across Exp. 2-4), indicating an overall bias towards confirmatory feedback.

**Supplementary Figure S6.**
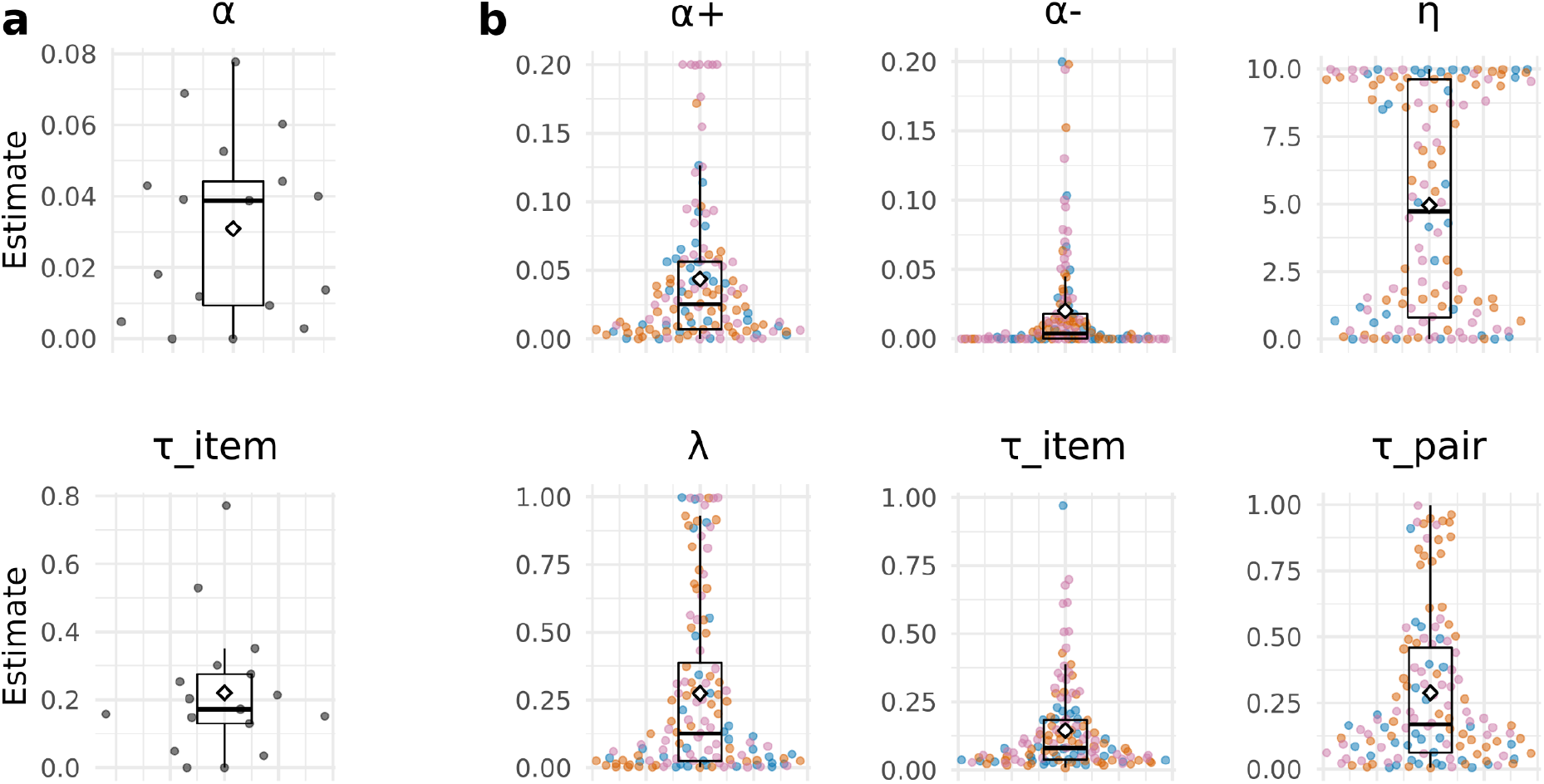
Individual parameter estimates for the winning model in each experiment (cf. Fig. 3). **a**, Experiment 1 (model Q1). **b**, Experiments 2-4 (model Q2*+P). Coloured dots show individual participant estimates, where blue, orange and pink colours refer to Experiment 2, 3 and 4, respectively. The diamond shape denotes the mean across experiments. While the estimates of *η* showed large variability with many values close to the upper bound, we observed no improvement in model fit when increasing the upper bound further.

**Supplementary Figure S7.**
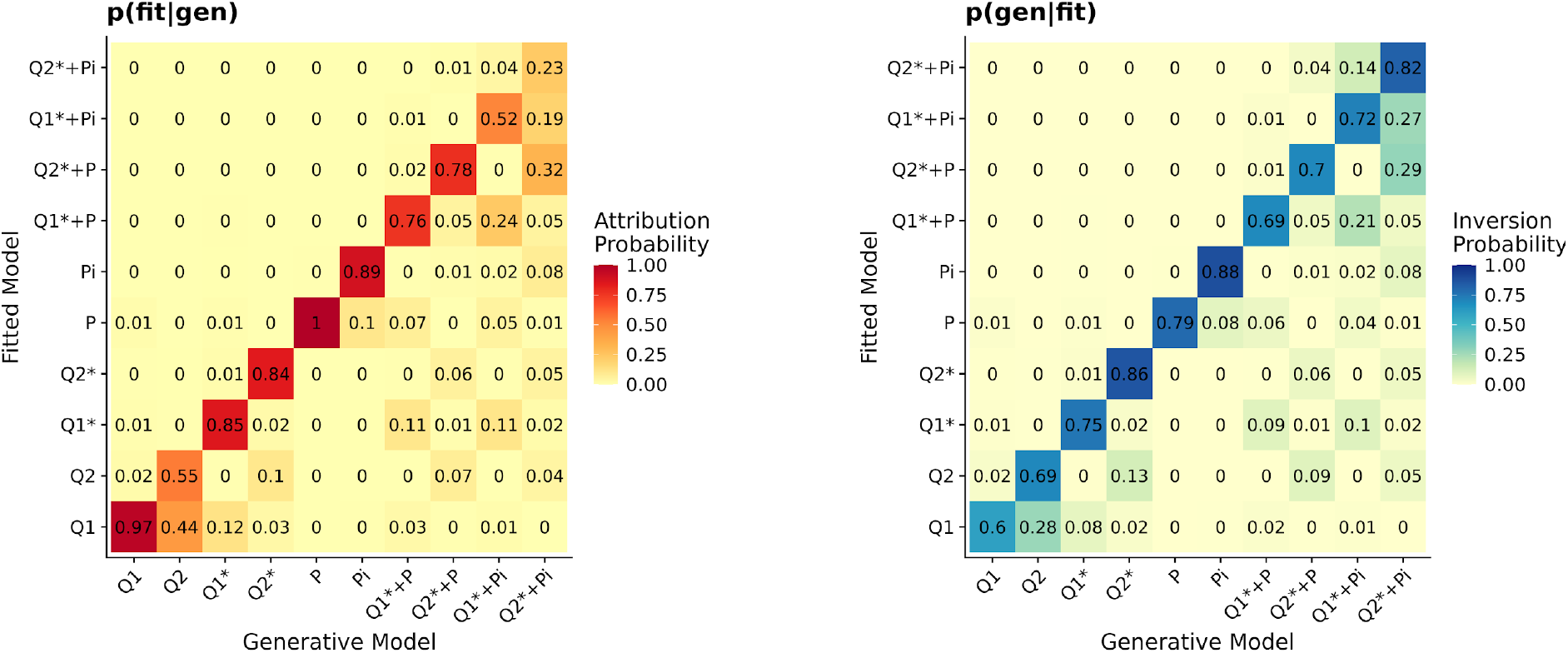
Model recovery results, partial feedback (cf. Fig. 3f). The models were generally well distinguished both in *p*(*fit*|*gen*) and in *p*(*gen*|*fit*). Of particular importance, our best asymmetric models (Q2* and Q2*+P, see *Results*) were well distinguished from their symmetric counterparts (Q1* and Q1*+P), with confusion rates no higher than 5%. Columns (left) and rows (right) may not always sum to 1 due to rounding of cell entries.

**Supplementary Figure S8.**
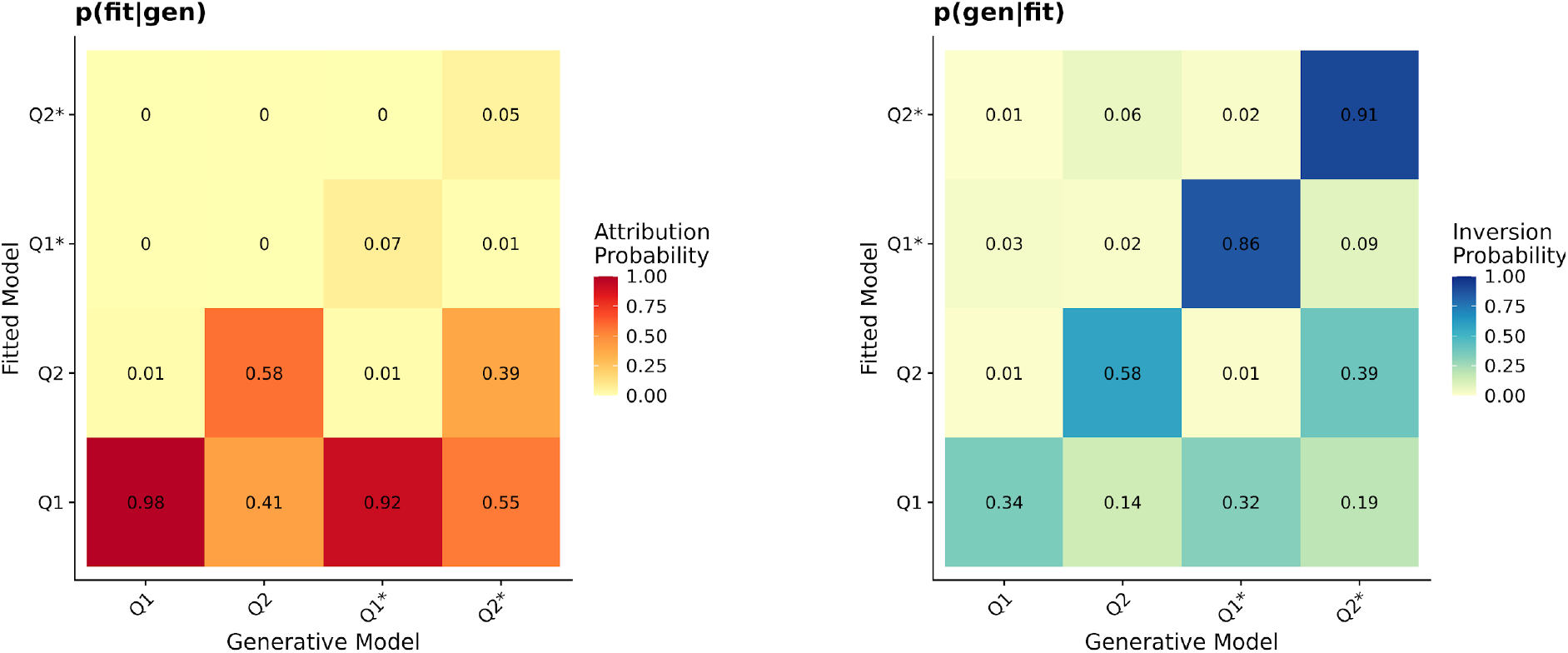
Model recovery results for Exp. 1 with full feedback (cf. Fig 3e). Simple Q-learning (models Q1/Q2) could not be confidently distinguished from models Q1*/Q2*. However, symmetric (Q1/Q1*) and asymmetric learning (Q2/Q2*) were distinguished relatively well. See *Methods*: *Model- and parameter recovery* for details.

**Supplementary Figure S9.**
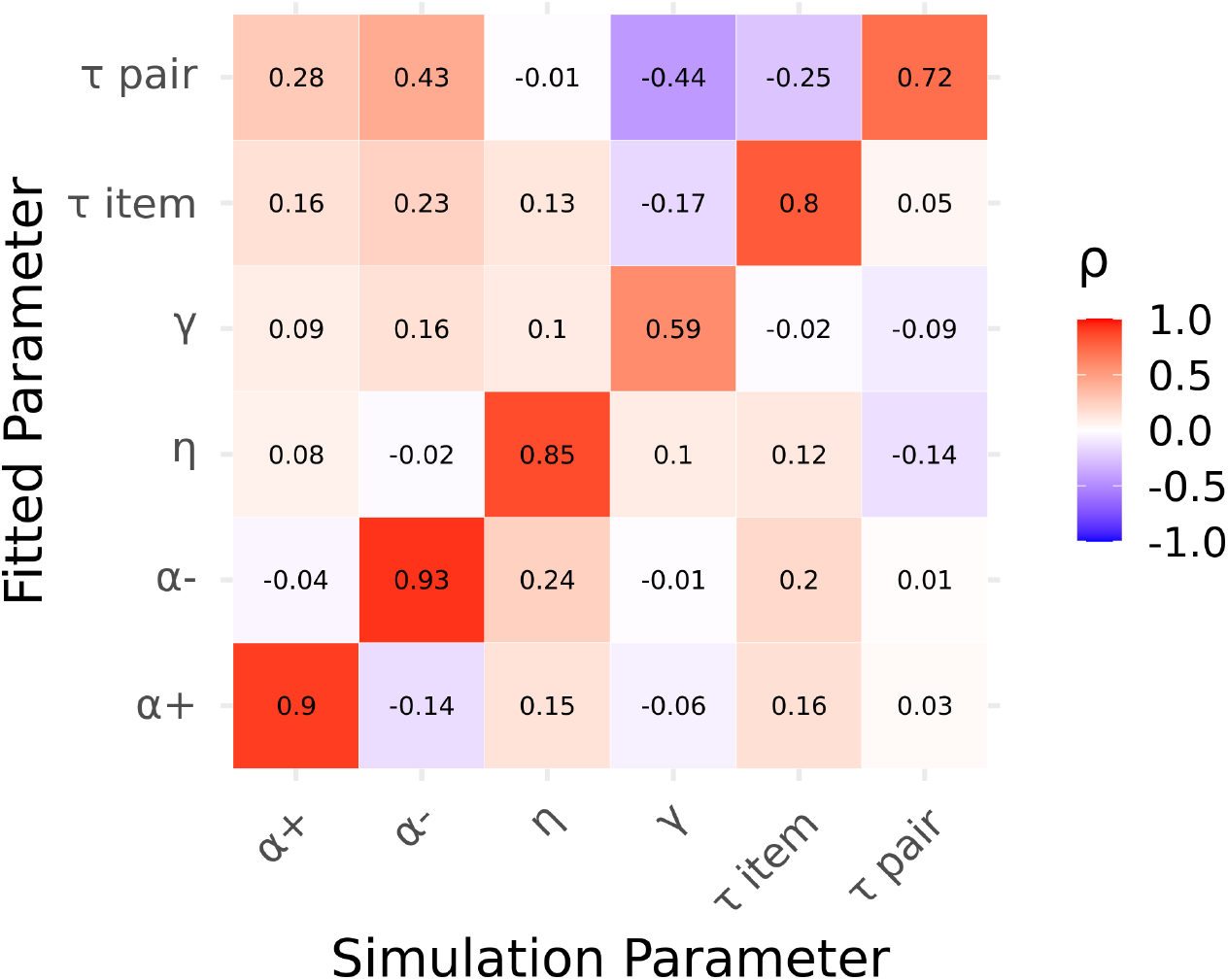
Parameter recovery results under partial feedback for our best-fitting model (Q2*+P). All fitted parameters correlate most strongly with their generative counterparts (diagonal) while correlations with other generative parameters (off-diagonal) are generally weaker.

**Supplementary Figure S10.**
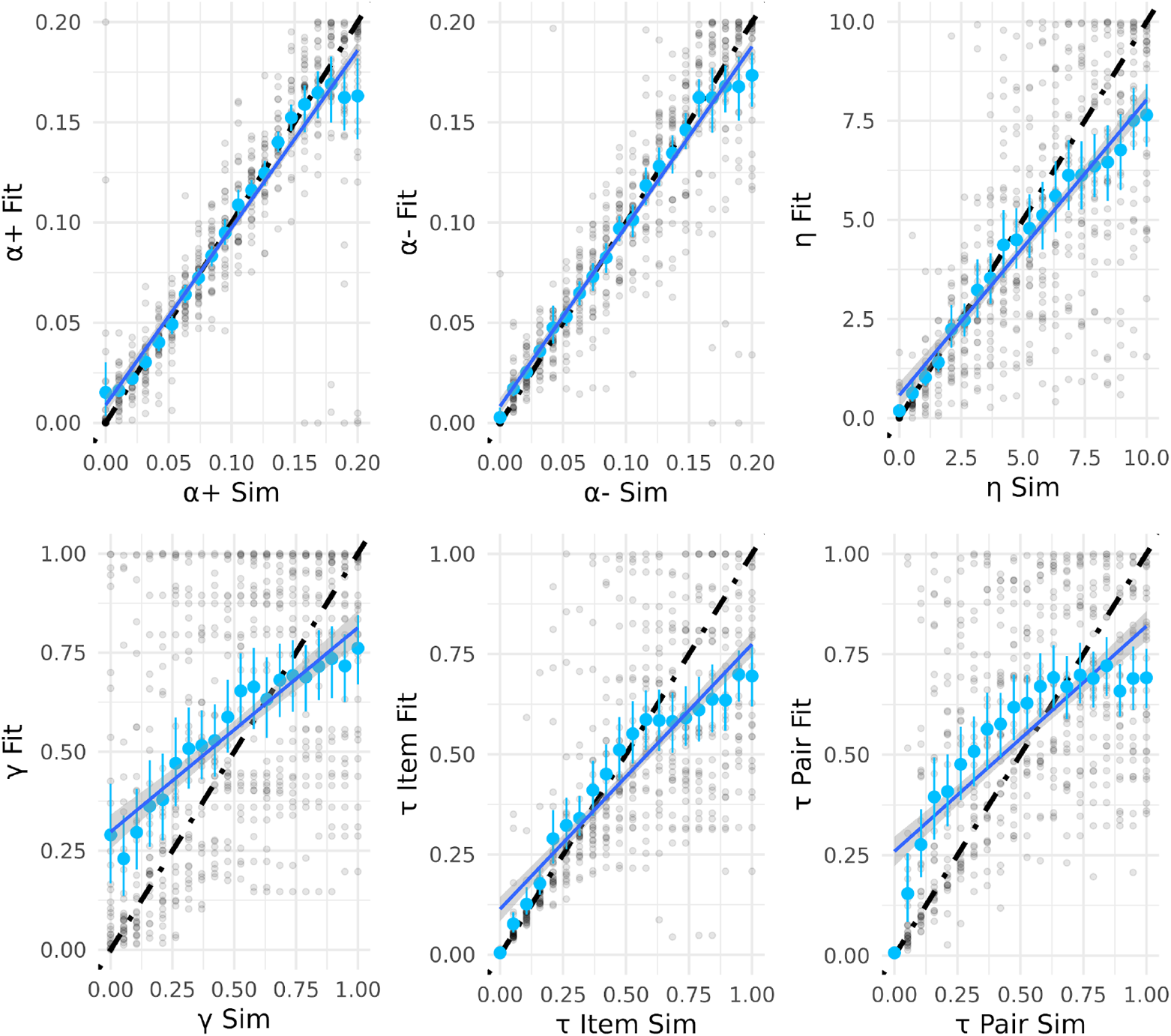
Detailed parameter recovery results for the individual parameters. The parameter values used to simulate choice data are plotted on the x-axes and the parameter estimates obtained from fitting the model to the simulated data are plotted on the y-axes. Light blue: mean recovered parameter values with bootstrapped 95% confidence intervals. Dark blue line shows linear fit. Results from individual recovery runs are shown as half-transparent black dots.

## Supplementary Methods

### RL-ELO

When fitting RL-ELO, we replaced our Q-learning process (*Methods*: *Item-level learning, Eq. 1*) by a rank learning process as proposed by Kumaran and colleagues^1^

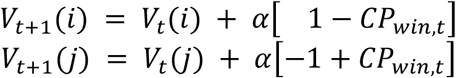

where *V*(*i*) and *V*(*j*) are the ranks of the winning item *i* and the losing item *j, CP*_*win*_ is the probability of choosing the winning item, and *α* is the learning rate. *CP*_*win*_ was computed with a logistic choice function (analogous to Eq. 5) of the difference in ranks between the winning and the losing item [*V*(*i*) − *V*(*j*)].

### Value-transfer

The value transfer model (VAT) proposed by von Fersen and colleagues^2^ assumes that the value of the losing item is updated with a proportion of the value of the winning item. We implemented VAT in a similar form as described previously^1^:

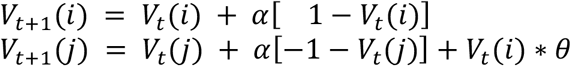

where *V*(*i*) and *V*(*j*) are the values of the winning item *i* and the losing item *j, α* is the learning rate, and *θ* controls the value transfer from the winning to the losing item. Interestingly, this formulation of VAT incorporates a form of asymmetric learning (through value transfer from winner to loser but not vice versa), and it can even predict below-chance performance for certain item pairings (through exceedingly large values of *θ*), similar to our Q2* model family. However, the Q2* process provided a better description of our empirical data (see *Results*).

For comparisons with our winning model (Q2*+P), we additionally fitted extended variants of RL-ELO and VAT where we included separate learning rates for winner and losers (*α*^+^ and *α*^−^, analogous to our model Q2, see *Methods*, equation 3) as well as pair-level learning (+P, equations 6-7 and 9-10).

